# Inert and seed-competent tau monomers suggest structural origins of aggregation

**DOI:** 10.1101/163394

**Authors:** Hilda Mirbaha, Dailu Chen, Olga A. Morozova, Kiersten M. Ruff, Apurwa Sharma, Xiaohua Liu, Rohit V. Pappu, David W. Colby, Hamid Mirzaei, Lukasz A. Joachimiak, Marc I. Diamond

**Affiliations:** Center for Alzheimer’s and Neurodegenerative Diseases, University of Texas, Southwestern Medical Center, Dallas, Texas 75390; Department of Chemical and Biomolecular Engineering, University of Delaware, Newark, Delaware 19716; Department of Biomedical Engineering, Washington University in St. Louis, St. Louis, Missouri 63130; Department of Biochemistry, University of Texas, Southwestern Medical Center, Dallas, Texas 75390

**Author notes:** Corresponding Author Marc I. Diamond, M.D. NL10.120, 5323 Harry Hines Blvd. Dallas, TX 75390.

## Abstract

Tauopathies feature progressive accumulation of tau amyloids. Pathology may begin when these amplify from a protein template, or seed, whose structure is unknown. We have purified and characterized distinct forms of tau monomer—inert (M_i_) and seed-competent (M_s_). Recombinant M_s_ triggered intracellular tau aggregation, induced tau fibrillization *in vitro*, and self-assembled. M_s_ from Alzheimer’s disease also seeded aggregation and self-assembled *in vitro* to form seed-competent multimers. We used crosslinking with mass spectrometry to probe structural differences in M_i_ vs. M_s_. Crosslinks informed models of local peptide structure within the repeat domain which suggest relative inaccessibility of residues that drive aggregation (VQIINK/VQIVYK) in M_i_, and exposure in M_s_. Limited proteolysis supported this idea. Although tau monomer has been considered to be natively unstructured, our findings belie this assumption and suggest that initiation of pathological aggregation could begin with conversion of tau monomer from an inert to a seed-competent form.

## Introduction

Amyloids are ordered protein assemblies, typically rich in beta sheet, that underlie multiple disorders such as Alzheimer’s disease (AD). Amyloid-forming proteins include tau, synuclein, and expanded polyglutamine proteins such as huntingtin, among many others. It is unknown how or why intracellular proteins such as tau transition from a relatively inert form to one that efficiently self-assembles into ordered structures *in vivo*. This process begins with the formation of a pathogenic “seed,” a structure that serves as a template for homotypic fibril growth. This structural transition could be a critical event in the pathogenesis of neurodegeneration. Under defined conditions and relatively high concentrations (typically micromolar), recombinant tau monomer will form amyloid fibrils *in vitro*. However the basis of spontaneous assembly in cells is unknown. The conversion of a protein from a monomer to a large, ordered multimer could occur by several mechanisms, but the first step probably involves the formation of a seed. This event, and indeed the actual conformation or assembly state of the protein that constitutes the “minimal” seed, has remained obscure. This has led to the idea that a seed is potentially transitory, arising from an equilibrium between two states: one relatively aggregation-resistant, and another that is short-lived. A seed could be a single molecule, or several. Based on extrapolation from kinetic aggregation studies, it has been suggested that a critical seed for tau and polyglutamine peptide amyloid formation is a single molecule^1, 2^, while an earlier study (among others^3^) has proposed a tau multimer^4^. Isolation of the seed-competent form of tau could be critical to understanding the initiation of disease and the design of more effective diagnostics and therapeutics.

Tau forms amyloids that underlie neurodegeneration in a variety of neuropathological syndromes, collectively termed tauopathies^5^. These include AD and frontotemporal dementias, among many others. Multiple groups, including ours, have now observed that tau will propagate an aggregated state from the outside to the inside of a cell, between cells, across synapses, and within brain networks^6^. In prior work we used size exclusion chromatography (SEC) to define tau trimers as the minimal unit of spontaneous cellular uptake and intracellular amyloid formation, and proposed this as the smallest particle capable of propagating aggregates between cells^7^. This work involved application of “naked” protein assemblies derived from recombinant protein or human brain onto cultured “biosensor” HEK293 cells or primary neurons that express a tau aggregation reporter^8, 9^. Biosensor cells and primary neurons alike take up tau aggregates via macropinocytosis^10^. The aggregates subsequently serve as highly specific templates to trigger intracellular amyloid formation^9, 11^. We have also determined that preincubation of cationic lipids such as Lipofectamine with tau seeds facilitates their direct transduction into a cell, bypassing the physiologic uptake mechanism^9, 12^. Lipofectamine-mediated delivery into biosensor cells allows direct quantitation of seed titer for both tau and *α*-synuclein^10^.

Tau is intrinsically disordered upon isolation from bacteria or mammalian cells and is relatively inert in terms of spontaneous self-assembly. However under various conditions, including exposure to polyanions such as heparin, tau will form aggregates via nucleated self-assembly ^13, 14^. It is unknown how these experimental conditions relate to the initiation of aggregation in human brain. We have now purified various stable forms of full-length tau monomer from recombinant sources and human brain. One is relatively inert and is stable for long periods. Another is “seed-competent,” triggers amyloid formation in cells and *in vitro*, and exhibits intrinsic properties of self-assembly. We have used crosslinking with mass spectrometry (XL-MS) to probe the structures of these molecules. Models of discrete regions within the RD predict that differential exposure of hexapeptide motifs previously known to be important for amyloid formation distinguishes the two forms of tau. These models are supported by limited proteolysis studies. The identification of distinct and stable forms of tau monomer, including some that are uniquely seed-competent, bears directly on how we understand the initiation of protein aggregation in the tauopathies.

## MATERIALS AND METHODS

### Tau expression, purification, fibrillization, and labeling

We utilized several forms of recombinant tau. Full-length (FL), wild-type (WT) tau contains two cysteines that create disulfide bridges and could complicate isolation of monomer. Thus in addition to preparing FL WT tau (2N4R) as previously described^15^, we purified FL tau (2N4R) that contains two cysteine/alanine substitutions (C291A, C322A), termed tau (2A). We used the 2A and WT forms of tau in our initial studies, before exclusively studying WT. Additionally, for fluorescence correlation spectroscopy (FCS), we engineered a single cysteine at the amino terminus (Cys-Tau (2A)) for labeling via maleimide chemistry. These modified proteins have fibrillization and seeding properties similar to FL WT tau. To initiate fibrillization, we incubated 8µM tau in 10mM HEPES, 100mM NaCl, and 8 µM heparin (1:1 ratio of FL tau to heparin) at 37°C for 72 h without agitation. For cysteine labeling, we incubated 200 µL of 8µM fibrils (monomer equivalent) and monomer with 0.025 mg of Alexa Fluor-488 (AF488) C5-maleimide (Invitrogen) and 80µM Tetramethylrhodamine-5-maleimide (Sigma-Aldrich) overnight at 4°C with gentle rotation. We quenched excess dye with 10mM DTT for 1h at room temperature. For limited heparin exposure, recombinant tau at 1µM was incubated with heparin at 1µM for 15min, 1hr and 4hr at 37°C before purification of monomer via Superdex 200 column.

To avoid confusion throughout the manuscript, we employ the following terminology:

**M_i_**: This refers to “inert” tau monomer, whether recombinant or derived from control brain.

**M_s_**: This refers to “seed competent” monomer, whether derived from sonicated fibrils, heparin-treated monomer, or AD brain.

### Sonication and size exclusion chromatography (SEC)

We sonicated labeled and non-labeled fibrils using a Q700 Sonicator (QSonica) at a power of 100-110 watt (Amplitude 50) at 4°C for 3h. Samples were then centrifuged at 10,000 x g for 10 min and 1 mL of supernatant was loaded into a Superdex 200 Increase 10/300 GL column (GE Healthcare) and eluted in PBS buffer at 4°C. After measuring the protein content of each fraction with a Micro BCA assay (Thermo Scientific) and/or fluorescence using a plate reader (Tecan M1000), we aliquoted and stored samples at −80°C or immediately used them in biochemical studies and cell seeding assays. Each aliquot was thawed immediately before use. The molecular weight/radius of proteins in each fraction was estimated by running gel filtration standards (Bio-Rad): Thyroglobulin (bovine) 670 kDa/845nm; γ-globulin (bovine) 158 kDa/5.29nm; Ovalbumin (chicken) 44 kDa/3.05nm; myoglobin (horse) 17 kDa/2.04nm; and vitamin B_12_ 1.35 kDa/0.85nm.

### Size-cutoff filtration

Monomer, dimer and trimer fractions were passed through a 100kDa MWCO filter (Corning) as instructed by the manufacturer (centrifuged at 15,000 x g for 15min at 4°C). Filtered material was immediately collected and used in seeding assay along with the non-filtered samples of the same fraction at a final concentration of 100nM, or analyzed by limited proteolysis. Protein concentration was determined before and after filtration by determining absorption at 205nm.

### CD spectroscopy

Circular dichroism (CD) measurements were performed at 25°C on a Jasco J-815 spectropolarimeter using a 0.1cm optical path length. 200μL of 2μM M_s_ or M_i_ monomer was dialyzed onto 10 mM NaP and the spectra were measured at 0.10 nm intervals, with a band width of 1.0nm, and scan speed of 10nm/min. The spectrum represents the average of 4 scans in the range of 195 to 250nm.

### Enzyme-linked immunosorbent assay

A total tau “sandwich” ELISA was performed similarly to that described previously^16^. Antibodies were kindly provided by Dr. Peter Davies (Albert Einstein College of Medicine). 96-well round-bottom plates (Corning) were coated for 48 hours at 4°C with DA-31 (aa 150-190) diluted in sodium bicarbonate buffer (6µg/mL). Plates were rinsed with PBS 3 times, blocked for 2h at room temperature with Starting Block (Pierce), and rinsed with PBS 5 additional times. SEC fractions were diluted in SuperBlock solution (Pierce; 20% SuperBlock, diluted in TBS), and 50 µL sample was added per well. DA-9 (aa 102-150) was conjugated to HRP using the Lighting-Link HRP Conjugation Kit (Innova Biosciences), diluted 1:50 in SuperBlock solution, and 50µL was added per well (15µg/mL). Sample + detection antibody complexes were incubated overnight at 4°C. Plates were washed with PBS 9 times with a 15 sec incubation between each wash, and 75 µL 1-Step Ultra TMB Substrate Solution (Pierce) was added. Plates were developed for 30min, and the reaction quenched with 2M sulfuric acid. Absorbance was measured at 450nm using an Epoch plate reader (BioTek). Each plate contained a standard curve, and all samples were run in triplicate.

### Fluorescence correlation spectroscopy

FCS measurements were conducted on a Confocal/Multiphoton Zeiss LSM780 Inverted microscope (Carl Zeiss-Evotec, Jena, Germany), using a 40X water immersion objective as previously described ^17^. Fluorescently labeled tau from SEC fractions (in PBS) was excited at 488nm and 561nm for 30sec, recording 10 times^18^. The data analysis was performed with Origin 7.0 (OriginLab, Northampton, MA).

### Liposome-mediated transduction of tau seeds

Stable cell lines were plated at a density of 35,000 cells per well in a 96-well plate. After 18h, at 60% confluency, cells were transduced with protein seeds. Transduction complexes were made by combining [8.75μL Opti-MEM (Gibco) +1.25μL Lipofectamine 2000 (Invitrogen)] with [Opti-MEM + proteopathic seeds] for a total volume of 20μL per well. Liposome preparations were incubated at room temperature for 20min before adding to cells. Cells were incubated with transduction complexes for 24h.

### FRET flow cytometry

Cells were harvested with 0.05% trypsin and fixed in 2% paraformaldehyde (Electron Microscopy Services) for 10min, then resuspended in flow cytometry buffer. The MACSQuant VYB (Miltenyi) was used to perform FRET flow cytometry. To measure CFP and FRET, cells were excited with a 405nm laser, and fluorescence was captured with 405/50nm and 525/50nm filters, respectively. To measure YFP, cells were excited with a 488nm laser and fluorescence was captured with a 525/50nm filter. To quantify FRET, we used a gating strategy similar to that previously described^9^. The integrated FRET density (IFD), defined as the percentage of FRET-positive cells multiplied by the median fluorescence intensity of FRET-positive cells, was used for all analyses. For each experiment, ∼20,000 cells were analyzed in triplicate. Analysis was performed using FlowJo v10 software (Treestar).

### Tau seeding *in vitro*

Recombinant full length (0N4R) tau monomer was purified as previously described^19^ at 1mg/mL in BRB80 buffer (80mM PIPES, 1mM MgCl2, 1mM EGTA, pH 6.8 with 0.3M NaCl) and boiled at 100°C for 5min with 25mM β-mercaptoethanol. The tau protein solution was then rapidly diluted 1:5 and cooled to 20°C in PBS, pH 7.4, to a final concentration of 0.2mg/mL of tau and 5mM β-mercaptoethanol. This solution was supplemented with Thioflavin T (ThT) to a final concentration of 20µM and filtered through a sterile 0.2µm filter. Reaction sizes of 195µL were aliquoted from the prepared protein stock and thoroughly mixed with 5µL of each sample at 100nM monomer equivalent, or 5µL of buffer control. For each sample, three different technical replicates were prepared. An opaque 96-well plate was prepared with a 3mm glass bead added to each well to increase agitation. The recombinant tau solution was added to the plate in 200µl reaction volumes. The plate was sealed with sealing tape to prevent evaporation and incubated in the plate reader (SpectraMax M2) at 37°C. ThT fluorescence was monitored over time with excitation and emission filters set to 444nm and 485nm, respectively. Fluorescence readings were taken every 5min, with agitation for 5sec before each reading.

### Tau extraction from brain and characterization by SEC

0.5g frontal lobe sections from AD patients at late Braak stage (VI) and age-matched controls lacking evident tau pathology were gently homogenized at 4°C in 5mL of TBS buffer containing protease inhibitor cocktails (Roche) using a dounce homogenizer. Samples were centrifuged at 21,000 x g for 15 min at 4°C to remove cellular debris. Supernatant was partitioned into aliquots, snap frozen and stored at −80°C. Immunopurification was performed with HJ8.5 anti-tau antibody^20^ at a ratio of 1:50 (1µg mAb per 50µg of total protein), incubating overnight at 4°C while rotating. To each 1mL of mAb/brain homogenate we added 200µL of a 50% slurry protein G-agarose beads (Santa-Cruz). We washed the bead with TBS buffer before overnight incubation at 4°C. We then centrifuged the complexes at 1000 x g for 3min and discarded the supernatant. Beads were washed with Ag/Ab Binding Buffer, pH 8.0 (Thermo Scientific) three times. Tau bound to the beads was eluted in 100 µL low pH elution buffer (Thermo Scientific), incubated at room temperature for 7min, followed by neutralization with 10µL Tris-base pH 8.5. This elution step was repeated once more with 50 µL elution buffer and 5µL Tris-base pH 8.5 for a total of 165µL. Samples were then centrifuged at 10,000 x g for 10min, and the supernatant loaded onto a Superdex 200 Increase 10/300 GL column (GE Healthcare). SEC fractions were frozen at −80°C after evaluation of protein content by Micro BCA assay (Thermo Scientific).

To compare different extraction methods, fresh frozen frontal lobe section from an AD patient brain was suspended in TBS buffer containing protease inhibitor cocktails (Roche) at 10% w/vol in 4 portions. Samples were homogenized using 3 different devices: a dounce homogenizer, probe sonicator (Omni International), and tissue homogenizer (Power Gen 125, Fischer Scientific). We also included one more condition of homogenizing with tissue homogenizer followed by probe sonication for 10min. Samples were centrifuged at 21,000 x g for 15min at 4°C to remove cellular debris. Supernatant was partitioned into aliquots followed by immunopurification.

To control for release of tau M_s_ from fibrils in AD brain, a tau KO Mouse brain was divided into two halves, followed by spiking one half with recombinant fibrils and the other with fibril-derived M_s_, both at final concentration of 10µM monomer equivalent. Each was dounce homogenized, centrifuged, immunoprecipitated with HJ8.5 anti-tau antibody, and fractionated by SEC with identical techniques as used for human brain processing. SEC fractions were then used in seeding experiments.

### Analysis of heat denaturation data

We analyzed the IFD from measurements of temperature dependent seeding using global fits to a proposed unimolecular heat denaturation reaction. This analysis rests on the Arrhenius equation^21^:

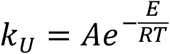

where *k_U_* is the unfolding rate constant, *E* is the activation energy, *R* is the gas constant, *T* is the temperature, and *A* is the pre-exponential factor. For the unimodal model, the data were fit globally to:

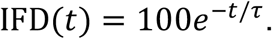

Here, *t* is the heat denaturation time and *τ* = 1/*k_U_* is the unfolding time. A second, multimodal model was deployed to account for discrepancies in the early time points which appeared to suggest the presence of a lag phase in denaturation. In this model, the data were fit globally to

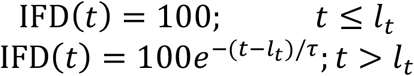

where *l*_2_ is the lag time given by

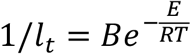

and *B* is a pre-exponential factor. We used the Akaike information criterion (AIC) to evaluate the best model as it quantifies the trade-off between goodness of fit and the complexity of the model ^22^. For least squares model fitting, AIC can be reduced to:

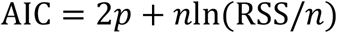

where *p* is the number of parameters in the model, *n* is the number of observations, and RSS is the residual sum of squares. The preferred model is the one with the minimum AIC. Here, we find AIC = 123 for the unimodal model and AIC = 105 for the multimodal model, which suggests the multimodal model is a better description of the denaturation data.

### Crosslinking, sample processing and LC-MS/MS analysis

M_i_ and M_s_ tau samples were prepared as described above. In all cases, tau preparations were crosslinked at a total protein concentration of ∼0.1mg/mL using 10 – 20µg starting material. The crosslinking buffer was 50 mM HEPES-KOH (pH 7.4) containing 150mM NaCl and 1mM DTT. The crosslinking reaction was initiated by adding disuccinimidyl suberate (DSS) stock solution (25 mM DSS-d_0_ and –d_12_, Creative Molecules) in DMF to a final concentration of 1mM. Samples were incubated at 37°C for 1min. For the heparin-derived M_s_ sample, heparin sulfate (Sigma) was added to a final concentration of 5µM, followed by 1mM DSS and the samples were incubated for 1min at 37°C. Excess reagent was quenched by addition of ammonium hydrogen carbonate to 50mM and incubation at 37°C for 30min, and then flash frozen at −80°C. Absence of higher molecular species was confirmed by SDS-PAGE and coomassie stain. After the quenching step, samples were evaporated to dryness in a vacuum centrifuge and resuspended in 8M urea. Proteins were reduced with 2.5mM TCEP (37°C, 30 min) and alkylated with 5mM iodoacetamide (30min, room temperature, protected from light). The sample solutions were diluted to 1M urea with 50mM ammonium hydrogen carbonate and trypsin (Promega) was added at an enzyme-to-substrate ratio of 1:50. Proteolysis was carried out at 37°C overnight followed by acidification with formic acid to 2% (v/v). Samples were then purified by solid-phase extraction using Sep-Pak tC18 cartridges (Waters) according to standard protocols. Samples were fractionated by size exclusion chromatography (SEC) on a Superdex Peptide column as described elsewhere ^23^. Two fractions collected from SEC were evaporated to dryness and reconstituted in water/acetonitrile/formic acid (95:5:0.1, v/v/v) to a final concentration of approximately 0.5 µg/µl. 2µL each were injected for duplicate LC-MS/MS analyses on an Eksigent 1D-NanoLC-Ultra HPLC system coupled to a Thermo Orbitrap Fusion Tribrid system. Peptides were separated on self-packed New Objective PicoFrit columns (11cm x 0.075mm I.D.) containing Magic C_18_ material (Michrom, 3µm particle size, 200Å pore size) at a flow rate of 300nL/min using the following gradient. 0-5min = 5 %B, 5-95min = 5-35 %B, 95-97min = 35-95 %B and 97-107min = 95 %B, where A = (water/acetonitrile/formic acid, 97:3:0.1) and B = (acetonitrile/water/formic acid, 97:3:0.1). The mass spectrometer was operated in data-dependent mode by selecting the five most abundant precursor ions (m/z 350-1600, charge state 3+ and above) from a preview scan and subjecting them to collision-induced dissociation (normalized collision energy = 35%, 30ms activation). Fragment ions were detected at low resolution in the linear ion trap. Dynamic exclusion was enabled (repeat count 1, exclusion duration 30sec).

### Analysis of mass spectrometry data

Thermo .raw files were converted into the open .mzXML format using msconvert (proteowizard.sourceforge.net) and analyzed using an in-house version of xQuest^24^. Spectral pairs with a precursor mass difference of 12.075321 Da were extracted and searched against the respective FASTA databases containing Tau (TAU_HUMAN P10636-8). xQuest settings were as follows: Maximum number of missed cleavages (excluding the crosslinking site) = 2, peptide length = 5-50 aa, fixed modifications = carbamidomethyl-Cys (mass shift = 57.021460 Da), mass shift of the light crosslinker = 138.068080 Da, mass shift of mono-links = 156.078644 and 155.096428 Da, MS^1^ tolerance = 10 ppm, MS^2^ tolerance = 0.2 Da for common ions and 0.3 Da for crosslink ions, search in ion-tag mode. For brain-derived samples we also included variable modifications including: Methionine oxidation = 15.99491, Ser/Thr/Tyr Phosphorylation = 79.96633 and Lysine Ubiquitylation = 114.043 with nvariable_mod = 1. Post-search manual validation and filtering of the recombinant samples was performed using the following criteria: xQuest score > 16, mass error between −4 and +7ppm, %TIC > 10, and a minimum peptide length of six aa. In addition, at least four assigned fragment ions (or at least three contiguous fragments) were required on each of the two peptides in a crosslink. False discovery rates (FDR) for the identified crosslinks were estimated using xprophet^24^. For the recombinant samples, M_i_ and M_s_, the FDR ranged from 6-10%. Post-search manual validation of the brain-derived samples was performed using the following criteria: xQuest score > 7, mass error between −5 and +7ppm, %TIC > 10, and a minimum peptide length of six aa. In addition, at least four assigned fragment ions (or at least three contiguous fragments) were required on each of the two peptides in a crosslink. The FDRs for the brain samples were much higher and ranged between 20-25%. For triplicate datasets corresponding to the M_i_ and M_s_ boiling time course we computed consensus crosslink profiles enforcing that at least two of the three datasets contain a crosslink. Crosslink data was visualized using Xvis^25^. Average contact distance was computed by averaging the sequence separation between crosslink pairs in a given dataset.

### Generation of structural models using XL-MS-derived constraints

High confidence crosslink pairs identified above were used to generate an ensemble of possible structures using a Rosetta protocol employing the crosslink pairs as structural restraints. The integration of XL-MS derived restraints have been previously used to refine structural models of large complexes^23^ and simpler heterodimeric complexes^26^. Based on distance distributions of crosslink pairs mapped onto crystallographic structures we set a lower bound of 15Å and an upper bound of 25Å for lysine C*α* pairs in our simulations. Importantly, in our simulations we weighted the constraint pairs as to allow some distances above the upper bound limit. The fragment library was supplanted by using chemical shifts derived from fibrillar tau ssNMR assignments (bmrb entry 17920) using csrosetta^27^. We generated 1000 models for each of the four XL-MS datasets on a high performance cluster (biohpc.swmed.edu). Representative structures were selected according to the low Rosetta score and radius of gyration. All plots were generated with gnuplot. All figures were generated using Pymol.

### Commandline used for *ab initio* protocol calculations with XL-MS restraints

AbinitioRelax.default.linuxgccrelease -in:file:fasta tau.fasta -file:frag3 tau.frags3.dat -file:frag9 tau.frags9.dat -nstruct 1000 -abinitio::increase_cycles 0.5 -abinitio::relax -score::weights score13_env_hb -abinitio::rg_reweight 0.5 -abinitio::rsd_wt_helix 0.5 -abinitio::rsd_wt_loop 0.5 -disable_co_filter true -out:file:silent csrosetta.out -constraints:cst_fa_file tau.cst - constraints:cst_file tau.cst -constraints:cst_weight 0.1 -constraints:cst_fa_weight 0.1 - loopfcst::coord_cst_weight 10.0

### Statistical analysis

Group mean values were analyzed by one-way ANOVA with Bonferroni post hoc significant differences test using GraphPad prism 5 software. Data in text and figures are represented as mean ± SEM.

### Kinetic analyses of M_i_ and M_s_ proteolysis

Limited proteolysis of M_i_/M_s_ using trypsin was carried out in 50mM TEAB at 25 °C. The enzyme to tau ratio was adjusted to 1:100 (wt/wt) with around 11ug of M_i_/M_s_ present initially. The total reaction mixture volume was 60µl. Aliquots were withdrawn from the reaction mixture at 1, 5, 15, 30, 60 and 120min by using 10µL of 10% trifluoroacetic acid (TFA) to quench the reaction (PH<3). The trypsin-digested peptides were then desalted using an Oasis HLB plate (Waters) and eluted with 100µL 80% acetonitrile (ACN) containing 0.1% TFA. The solvent was evaporated in a SpeedVac concentrator and the dried samples were reconstituted in 20µl of 2% acetonitrile, 0.1% TFA and 2µl solution was used for by LC/MS/MS analysis, the analysis were performed on an Orbitrap Elite mass spectrometer (Thermo Electron) coupled to an Ultimate 3000 RSLC-Nano liquid chromatography systems (Dionex). Samples were injected onto a 75µm i.d., 15-cm long EasySpray column (Thermo), and eluted with a gradient from 1-28% buffer B over 60 min. Buffer A contained 2% (v/v) ACN and 0.1% formic acid in water, and buffer B contained 80% (v/v) ACN, 10% (v/v) trifluoroethanol, and 0.1% formic acid in water. The mass spectrometer operated in positive ion mode with a source voltage of 2.8kV and an ion transfer tube temperature of 275 °C. MS scans were acquired at 240,000 resolution in the Orbitrap and up to 14 MS/MS spectra were obtained in the ion trap for each full spectrum acquired using collision-induced dissociation (CID), with charge 1 ions rejected. Dynamic exclusion was set for 15s after an ion was selected for fragmentation. Raw MS data files were searched against the appropriate protein database from Uniprot, and reversed decoy sequences appended (Elias and Gygi, 2007) by using Protein Discovery 2.2 (Thermo Fisher Scientific). Fragment and precursor tolerances of 20ppm and 0.6Da were specified, and 12 missed cleavages were allowed.

Carbamidomethylation of Cys was set as a fixed modification and oxidation of Met was set as a variable modification. Label-free quantitation of proteins across samples was performed. Average peptide intensity values were computed for all time points for each peptide. To estimate differences in kinetic profiles we calculated the median value of each profile and compared the M_i_ to M_s_ ratio.

## RESULTS

### Isolation of fibril-derived monomer and other assemblies

We initially sought to define the tau seeding unit that would trigger intracellular aggregation upon direct delivery to the cell interior. We had previously observed that a tau trimer is the minimal assembly size that triggers endocytosis and intracellular seeding^7^. These experiments depended on spontaneous cell uptake, since no Lipofectamine was added to the reactions. A prior study had also indicated the role of disulfide linkages in promoting tau aggregation, potentially by dimer formation^4^. Thus, for our initial studies we engineered and purified full-length (FL) tau monomer that lacks any internal cysteines due to alanine substitutions (C299A and C322A), termed tau (2A). FL tau (2A) cannot self-associate based on disulfide linkages, which helped prevent the formation of cryptic dimers that could have confounded our studies. These substitutions did not affect tau purification, heparin-induced fibrillization, and sonication protocols, which we performed as described previously^7^. We covalently labeled the fibril preps prior to sonication and isolation of recombinant FL tau (2A) assemblies of various sizes by size exclusion chromatography (SEC)^7^. In parallel, we also studied FL wild type (WT) tau. We purified unfibrillized recombinant FL tau (2A) monomer by SEC (Fig. 1A), and isolated SEC fractions of sonicated fibrils that contained putative monomer, dimer, trimer and ∼10-mer (Fig. 1B).

**Figure 1:**
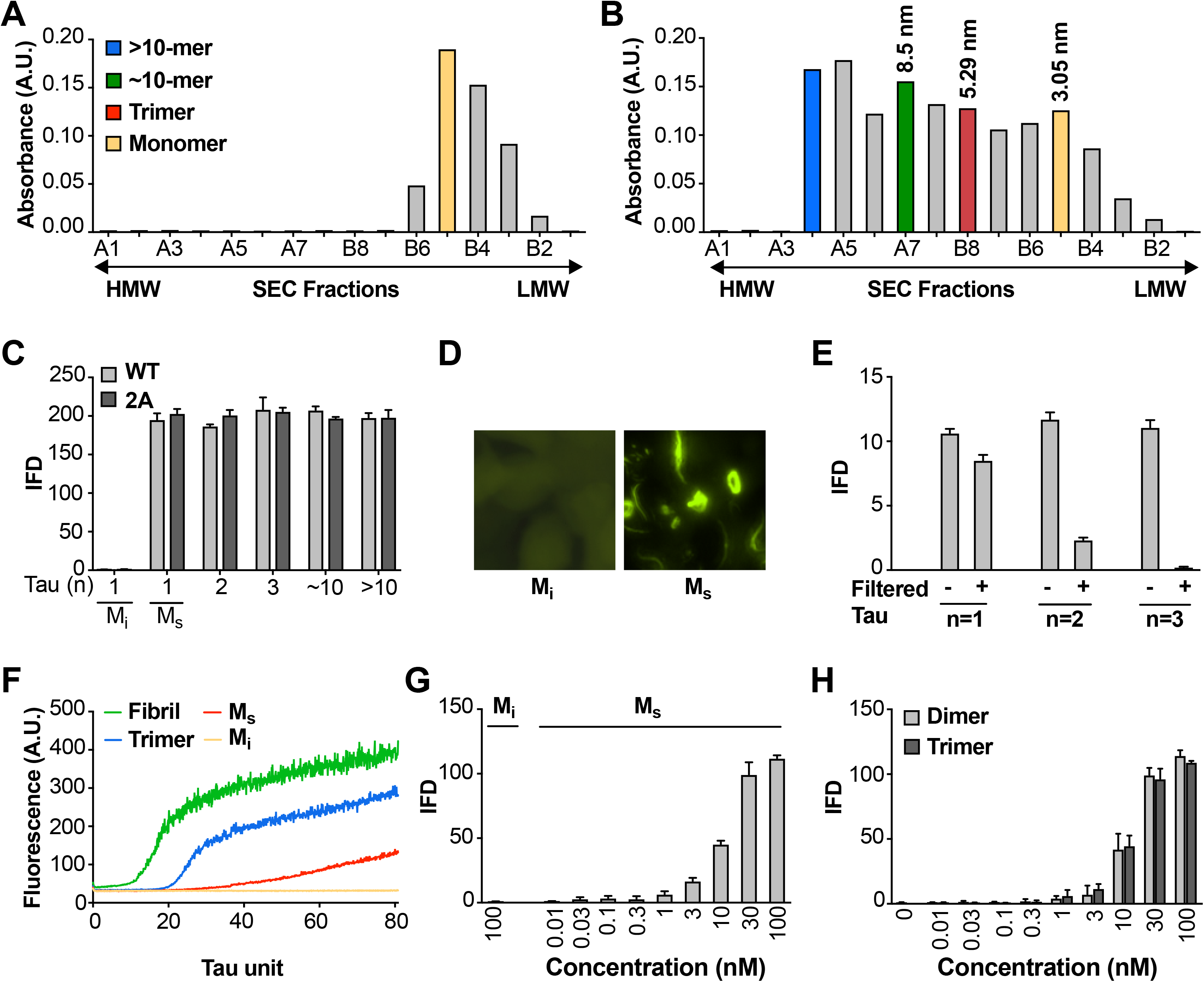
Seeding activity of tau monomer in cells and *in vitro*. (**A, B**) FL Cys-Tau(2A) was labeled with Alexa488 and resolved by SEC (**A**), or was fibrillized in the presence of heparin, labeled with Alexa488, sonicated, and the assemblies resolved by SEC (**B**). The column was calibrated using standards of the indicated hydrodynamic radii. Color codes indicate the putative assembly units. (**C**) Tau assemblies were seeded into tau RD-CFP/YFP biosensor cells. M_i_ represents “inert” monomer purified in (A), which had no seeding activity; M_s_ represents “seed-competent” monomer purified in (B), which induced intracellular tau aggregation. (**D**) FL WT tau and FL Cys-Tau(2A) were similarly fibrillized, sonicated, and the fragments resolved by SEC. Seeding activity of each fraction was determined. M_s_ and larger assemblies of both forms of tau exhibited seeding activity, but not M_i_. IFD = Integrated FRET Density. (**E**) Tau assemblies of n=1,2,3 were passed through a 100kD size cutoff filter. Filtration had no effect on the M_s_ fraction, whereas it reduced seeding of assemblies of n=2 or 3. (**F**) Tau fibrils, trimer, or monomer were used to induce fibrillization *in vitro* of full-length (0N4R) tau, measured by induced thioflavin fluorescence. M_i_ had no seeding activity, whereas M_s_, trimer, and unfractionated fibrils had strong seeding activity. (**G,H**) Titration of assemblies was performed. (**G**) M_s_ exhibited an EC_50_ of approximately 10nM (monomer equivalent); (**H**) Dimer and trimer had similar potencies. Concentration is reflected as monomer equivalent.

### Fibril-derived monomer exhibits seeding activity in cells and *in vitro*

To test the seeding activity of the tau preparations, we used a previously described “biosensor” cell reporter line^9^. These cells stably express 4R tau repeat domain (RD) containing the disease-associated P301S mutation, fused to cyan and yellow fluorescent proteins (RD-CFP/YFP). Exogenously applied seeds induce intracellular aggregation with resultant fluorescence resonance energy transfer (FRET) that can be measured via flow cytometry^9, 12^. The degree of aggregation is scored using “integrated FRET density” (IFD), which is the product of the percent positive cells and the mean fluorescence intensity of FRET-positive cells, and from this we determine a titer of tau seeding activity^9^. Lipofectamine directly transduces tau assemblies across the plasma membrane and increases the assay’s sensitivity by approximately 100-fold. Upon incubation with Lipofectamine, we were surprised to observe seeding by monomer and larger assemblies alike, whether FL WT or 2A. (Fig. 1C,D). Epifluorescence microscopy confirmed the presence of intracellular inclusions after FL WT tau monomer seeding (Fig. 1D). We termed the inert monomer “M_i_,” and the seed-competent monomer “M_s_.” To rule out higher order assemblies of tau within the putative monomer fraction, immediately prior to the seeding assay we passed fractions through a 100kDa cutoff filter to eliminate anything larger than a monomer. While monomer fraction retained ∼80% of seeding activity, only ∼20% of dimer seeding activity remained, and ∼1-2% of trimer seeding activity remained (Fig. 1E). To exclude an artifact related to Lipofectamine transduction into cells, we tested FL (2A) tau preparations in an *in vitro* seeding assay that induces fibril formation by full-length tau (0N4R) through iterative polymerization and agitation steps^19^. M_i_ had no intrinsic seeding activity. However M_s_ induced amyloid formation, albeit more slowly than trimer or unfractionated fibrils (Fig. 1F). This slow aggregation process may reflect inefficient fibril assembly, and a predominance of small nucleated assembly events from the added monomer. We concluded that the M_s_ fraction contained seeding activity that enabled intracellular aggregation of tau RD-CFP/YFP in cells, or full-length tau *in vitro*. Finally, we tested whether contamination of very small amounts of seeds could somehow account for the seeding activity in monomer fractions by carrying out dose-response titrations of the various preparations. M_s_ had an EC_50_ of ∼10nM (Fig. 1G), which was very similar to dimer and trimer (Fig. 1H). Thus to account for signal observed in the seeding assay, contamination of an otherwise inert monomer with larger seed-competent assemblies would have to be substantial.

### Comparison of M_i_ and M_s_ by CD and FCS

We tested for obvious structural differences between M_i_ and M_s_ using CD spectroscopy, which revealed none (Fig. 2A). We re-tested the assemblies using fluorescence correlation spectroscopy (FCS), which measures particle diffusion through a fixed volume. As we previously observed^7^, we accurately estimated the units of small assemblies (≤10-mer), but not larger assemblies (>10-mer) (Fig. 2B). In an additional effort to detect cryptic multimers within the M_s_ preparation, we used double-label FCS. We engineered a cysteine onto the amino terminus of FL tau (2A) to enable its covalent modification (Cys-Tau (2A)). We then prepared Cys-tau (2A) fibrils, or monomer, and labeled them simultaneously with Alexa488 (green) and tetramethylrhodamine (TMR) via maleimide chemistry. We carried out sonication and purification by SEC as before, isolating assemblies of various sizes. We evaluated each for cross-correlation between red and green signal, which indicates the presence of at least two tau molecules in a particle. We analyzed >300 events for each assembly. When we evaluated M_i_ and M_s_, 100% of events in each case showed a diffusion time consistent with a tau monomer (Fig. 2C,D). Furthermore, we observed no cross-correlation between red and green signal, indicating that neither preparation had detectable multimeric assemblies (Fig. 2C,D,H). By contrast, when we evaluated larger species such as dimer, trimer, or ∼10-mer, we observed longer diffusion times consistent with the predicted assembly sizes, and significant cross-correlation values (Fig. 2E-H), consistent with the presence of multimers. The FCS studies supported the conclusion that M_i_ and M_s_ are comprised predominantly of monomer.

**Figure 2:**
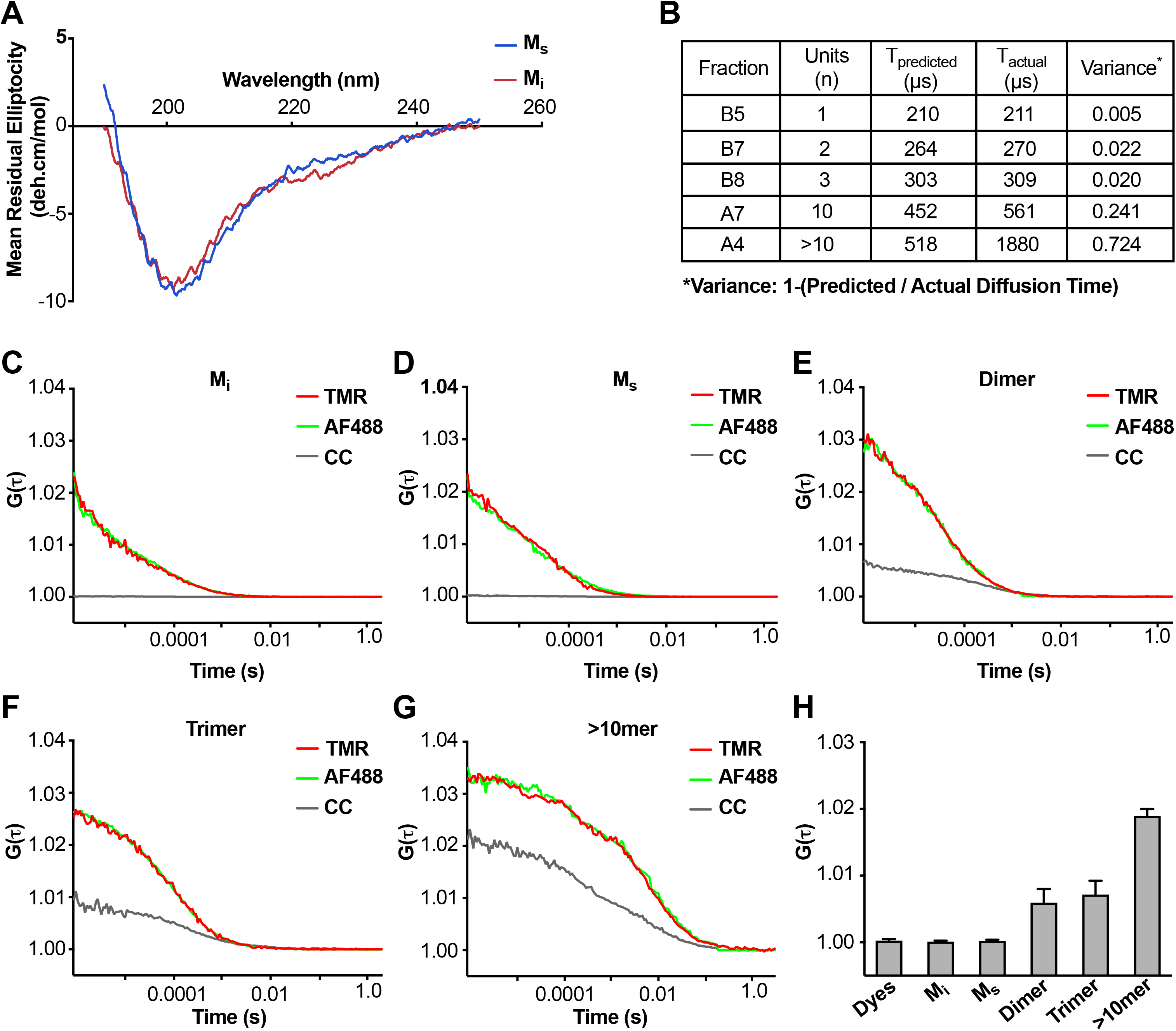
Analyses of M_i_ and M_s_ by CD and FCS. (**A**) CD spectra of M_i_ and M_s_ were similar. (**B**) FCS Diffusion times for M_i_, M_S_, dimer, trimer, and ∼10mer, and the cross-correlation for M_i_, M_s_, dimer, trimer, and ≥10-mer were determined after labeling of fibrils with Alexa488, or double labeling additionally with tetramethylrhodamine prior to sonication. Table reflects the predicted diffusion time and the actual diffusion time. The variance between predicted vs. observed times is reported. (**C-G**) FCS for double-labeled tau assemblies. Cross correlation (CC) between the two dyes is indicated in grey lines. (**H**) Summary of FCS cross-correlation, including free dyes. Neither free dye, M_i_ nor M_s_ showed any cross-correlation, indicating that single species predominate. All multimeric assemblies exhibited cross-correlation, indicating detection of both dyes within a single particle.

### SEC preparation efficiently purifies M_s_ monomer

To rule out cross-contamination of assemblies within the SEC column, we tested its ability to exclude larger seeds from the monomer fraction. We first isolated M_s_ and larger assemblies from a sonicated fibril preparation (Fig. 3, Group 1). Removing the fraction that contained M_s_ (B5), we then pooled the remaining fractions, and spiked them with M_i_. We re-fractionated the material on SEC to isolate the monomer in fraction B5 again (Fig. 3, Group 2). As previously observed, M_s_ and other fibril-derived assemblies in Group 1 had seeding activity (Fig. 3). However, in Group 2, while we observed seeding activity in larger assemblies, the monomer (which we take to be M_i_) re-isolated from a pool of larger fibril-derived assemblies had no seeding activity (Fig. 3). This confirmed that larger, seed-competent assemblies do not appreciably contaminate the monomer fraction during SEC.

**Figure 3:**
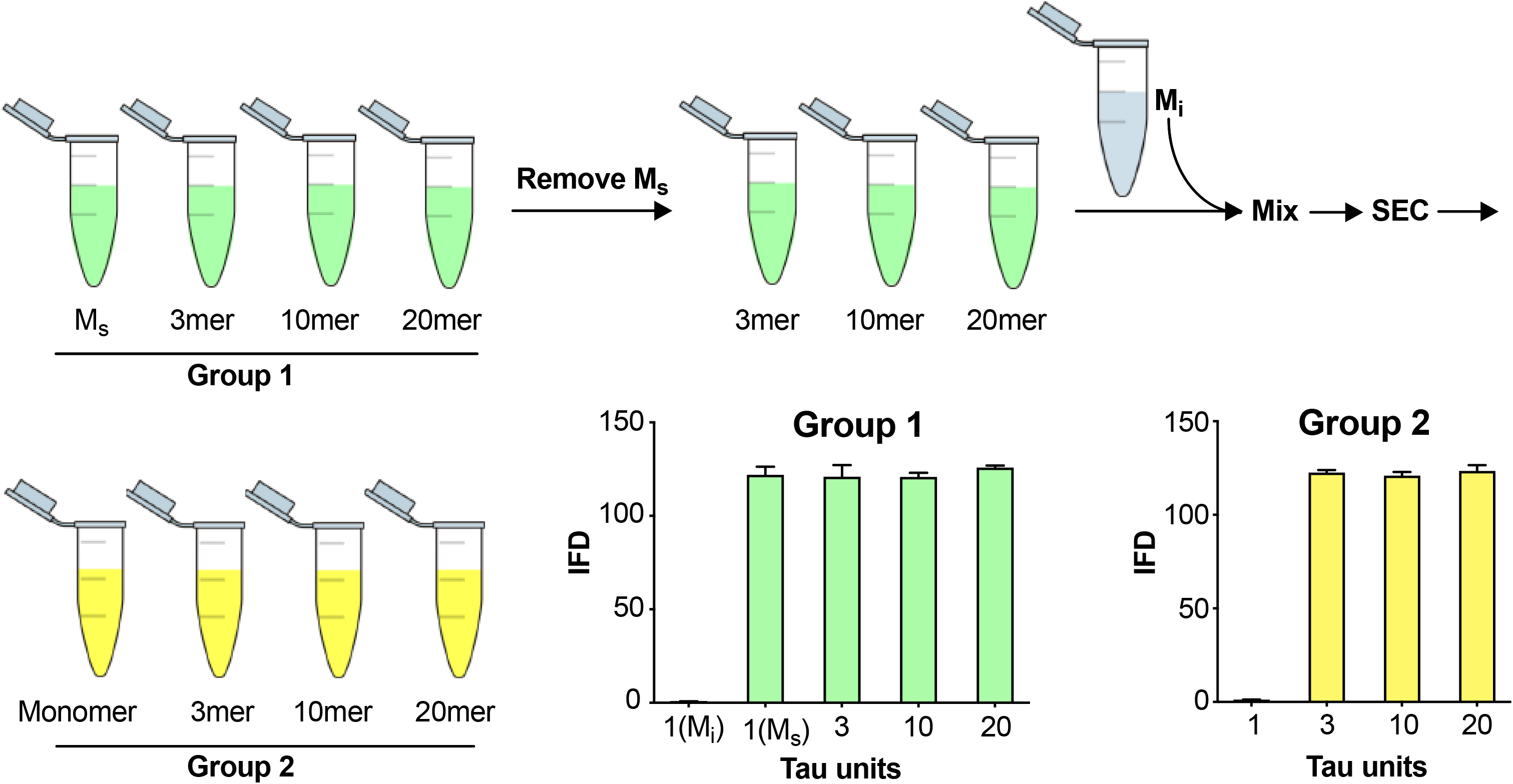
Fidelity of SEC purification of assemblies. SEC fidelity was tested by isolating M_s_ from fractions after fibril sonication. Remaining fractions were combined with M_i_, and the mix was re-isolated by SEC. In Group 1, after the first isolation, the monomer fraction (which contains M_s_) contained seeding activity. In Group 2, after the second purification by SEC, the monomer fraction (which contains M_i_ spiked in) did not exhibit seeding activity.

### Heat denaturation of assemblies

Although prior controls had essentially excluded the presence of tau multimers in the sample, we used heat-mediated dissociation of oligomeric assemblies as an additional test for the possibility that M_s_ in fact represents a uniquely compact multimer that somehow purifies as a monomer. We collected M_s_ by SEC, and heated the sample to 95°C for 3h. We then re-isolated the sample via SEC. We carried out the same procedure with trimer and ∼20-mer. In each case, we tested the resultant fractions for seeding activity. In the first instance, after heating we re-isolated M_s_ purely as monomer that retained virtually all of its seeding activity (Fig. 4A). The trimer assembly (fraction B8) broke down to smaller assemblies, predominantly monomer, each of which retained seeding activity (Fig. 4B). The ∼20-mer (fraction A5) was largely stable following heat treatment, and retained its seeding activity (Fig. 4C). These experiments highlighted the lability of small multimers (i.e. trimer), and a surprising persistence of seeding activity in heat-treated monomer.

**Figure 4:**
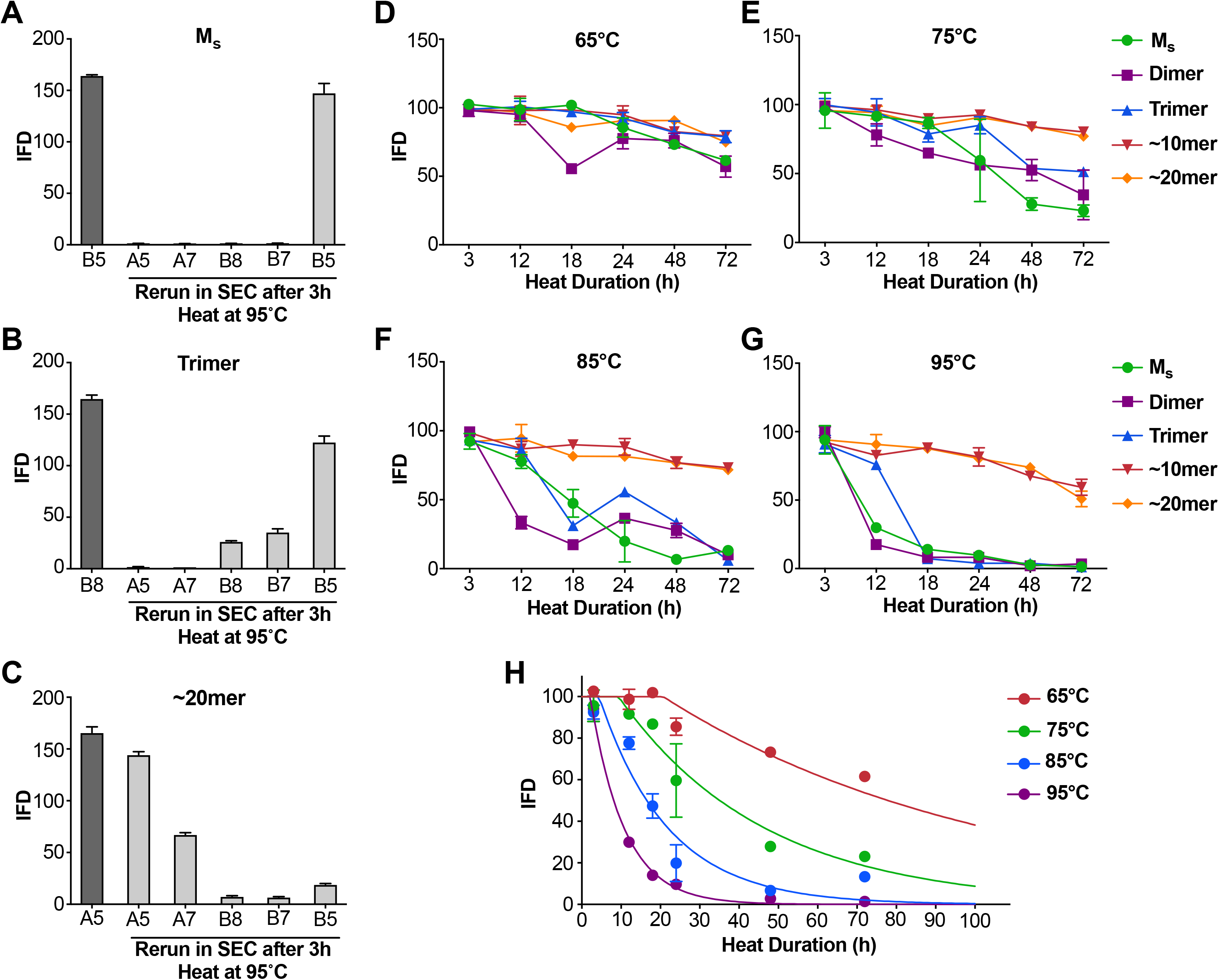
Heat denaturation of assemblies. (**A-C**) Heat-induced dissociation of assemblies. (**A**) The SEC fraction containing M_s_ (B5) was heated to 95°C for 3h and re-isolated by SEC prior to testing the FRET biosensor assay. No loss in seeding activity was observed. (**B**) When the SEC fraction containing trimer (B8) was heated similarly, seeding activity shifted to fractions that contain dimer and monomer (B7, B5). (**C**) ∼20-mer (A5) was largely stable to heating, although some smaller seed-competent assemblies were liberated. (**D-G**) Various assemblies were subjected to heat denaturation at the indicated temperatures and times, followed by analysis of seeding activity in the FRET biosensor assay. Whereas ∼10-mer and ∼20-mer were relatively stable from 65-95°C, monomer, dimer and trimer showed temperature-dependent loss of seeding activity. (**H**) Plot of denaturation data for M_s_ with multimodal regression curves superimposed.

### Differential heat lability of tau assemblies

In the preceding experiment M_s_ retained seeding activity even after 3h at 95°C, a condition sufficient to dissociate trimers. These experiments implied that M_s_ consists of a stable seed-competent structure, resistant to heat denaturation. Consequently, we used more nuanced heat denaturation of seeding activity to probe the relative stabilities of M_s_, dimer, trimer, and larger assemblies of FL WT tau. We first isolated tau monomer, dimer, trimer, ∼10-mer, and ∼20-mer on SEC. We then incubated the various assemblies at a range of temperatures (65, 75, 85, 95°C) and times (0, 3, 12, 18, 24, 48, 72h) before measuring seeding activity. Lower temperatures only slightly reduced seeding activity, whereas exposure of M_s_, dimer, and trimer to temperatures ≥85°C for 18-24h eliminated it at roughly the same rate for each (Fig. 4D-G). By contrast, the seeding activities of ∼10-mer and ∼20-mer were relatively heat-resistant (Fig. 4D-G). This was consistent with our prior observations that tau seeds derived from cultured cells are resistant to boiling ^11^. To determine a putative energy barrier between M_s_ and M_i_, we evaluated the denaturation data for M_s_ by integrating the data from the prior experiments (Fig. 4H). We compared two models for the transition of M_s_ to an inert form (which we assumed to be an unfolding reaction): a unimodal unfolding model vs. a multimodal model that assumes intermediate seed-competent states. The unimodal model did not account for the data at early time points, which indicated a lag phase in denaturation, whereas the multimodel model performed better (Fig. 4H). The lag phase in denaturation implied an ensemble of seed-competent states that define M_s_, each separated by smaller energy barriers. Using the multimodal model, we calculated the barrier to conversion of M_s_ to an inert form to be ∼78 kcal/mol.

### M_s_ has unique properties of self-assembly

Aggregation of M_i_ *in vitro* is relatively slow, requires high protein concentration (micromolar), and polyanions such as heparin^13, 14^. Based on the seeding activity of M_s_ we predicted that it might more readily self-associate. We incubated FL WT tau M_i_ and M_s_ alone, or dimer or trimer at equimolar ratios, keeping total particle concentration constant at 500nM. We then monitored change in assembly size over 24h. M_i_, dimer, and trimer showed no evidence of self-association in this timeframe (Fig. 5A,C,D). By contrast, when incubated alone, M_s_ readily formed larger assemblies (Fig. 5B). When we incubated M_i_ with dimer or trimer, we saw no change in the assembly population over 24h (Fig. 5E,F). By contrast, when we mixed M_s_ with dimer or trimer we observed a growth of larger assemblies with a concomitant reduction in dimer and trimer peaks (Fig. 5G,H). We conclude that M_i_, dimer, and trimer do not form larger assemblies at an appreciable rate, while M_s_ self-assembles and adds on to larger assemblies.

**Figure 5:**
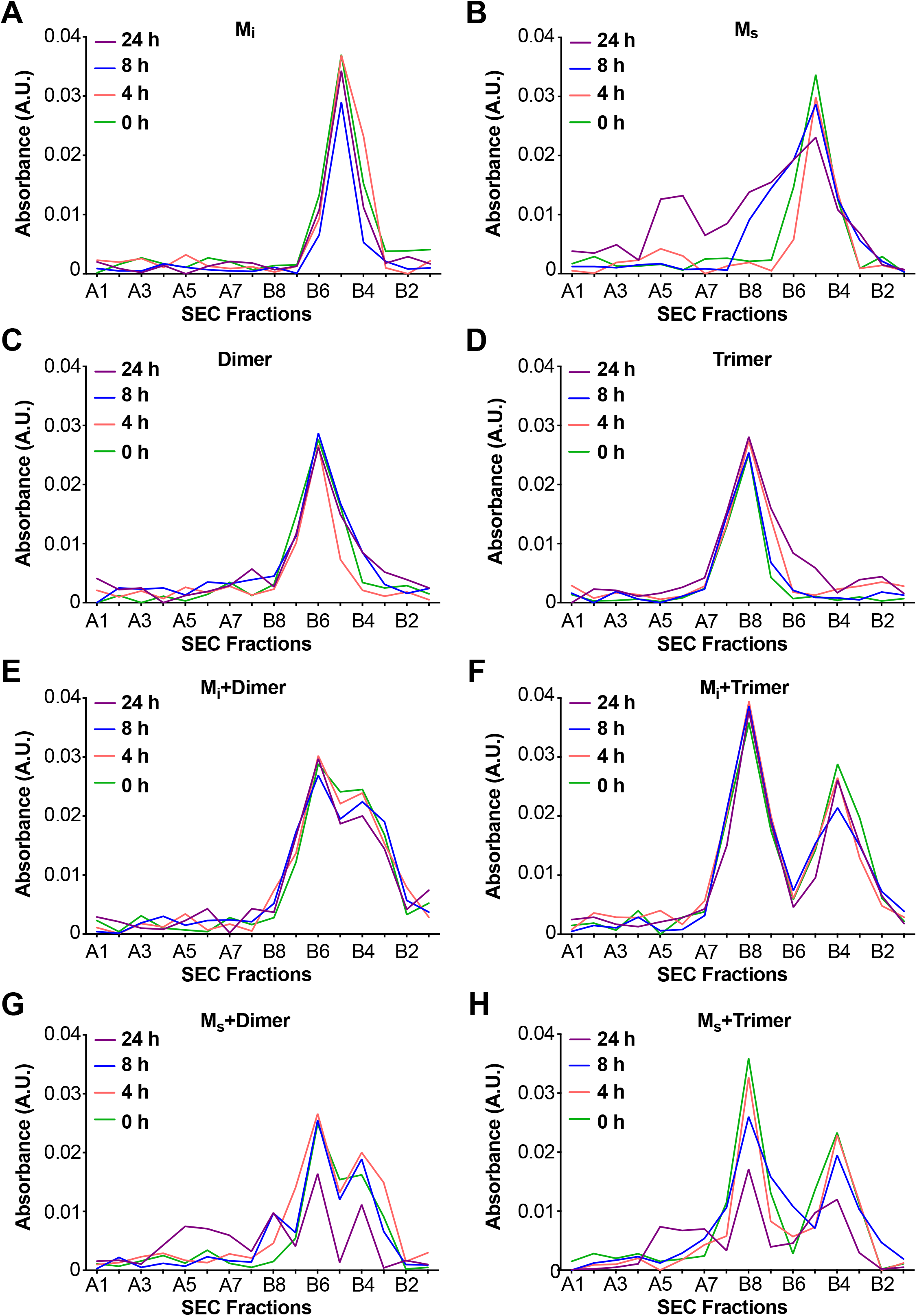
M_s_ self-assembles. M_i_ and M_s_ were incubated at 500nM or with equivalent amounts (monomer equivalent) of dimer and trimer for various times prior to resolution by SEC. Assemblies were monitored by reading the absorbance of fractions using micro BCA assay. (**A**) M_i_ showed no self-association. (**B**) M_s_ exhibited self-association over time. (**C,D**) Dimer and trimer were stable over time. (**E,F**) M_i_ does not react with dimer or trimer to form larger assemblies. (**G,H**) M_s_ reacts with dimer and trimer to form larger assemblies.

### Heparin induces transition from M_i_ to M_s_

The preparation of M_s_ based on sonication of fibrils raised two important issues. First, it left uncertain whether M_i_ could be converted to a seed-competent form without previously being incorporated into a fibril. Second, we observed that sonication could create fragments from tau monomer that might potentially act as seeds (Supp. Fig. S6A). Consequently, we used heparin to induce the formation of M_s_, thereby avoiding sonication. We exposed FL WT tau to heparin for varying amounts of time before purifying different assembly sizes by SEC and testing for seeding activity. After 15min of heparin exposure, we detected low but significant amounts of seed-competent monomer, and much fewer larger assemblies (Fig. 6A). Crosslinking of purified, heparin-induced M_s_ revealed no evidence of multimers or an increase in fragments (Supp. Fig. S6B). Recombinant monomer not treated with heparin had no seeding activity at any time point (Fig. 6A). At longer heparin treatment times (1h, 4h) monomer fractions as well as larger assemblies all had strong seeding activity (Fig. 6A). M_s_ derived from heparin exposure was relatively resistant to heat denaturation at 95°C, albeit less so than fibril-derived M_s_ (Fig. 6B). Relative seeding efficiency of the various forms of M_s_ as well as sonicated or unsonicated fibrils were relatively similar (Fig. 6C). We noted also that sonication of M_i_ and purification by SEC did not produce any seed-competent species, eliminating the possibility that small assemblies of sonication-induced fragments accounted for seeding activity of M_s_ (Fig. 6C). These experiments also indicated that it is not necessary for tau monomer to be part of a fibril or to be exposed to sonication to produce an efficient seed-competent monomer. Heparin, presumably by catalyzing a transition from an inert to a seed-competent form, enables this critical conformational change.

**Figure 6:**
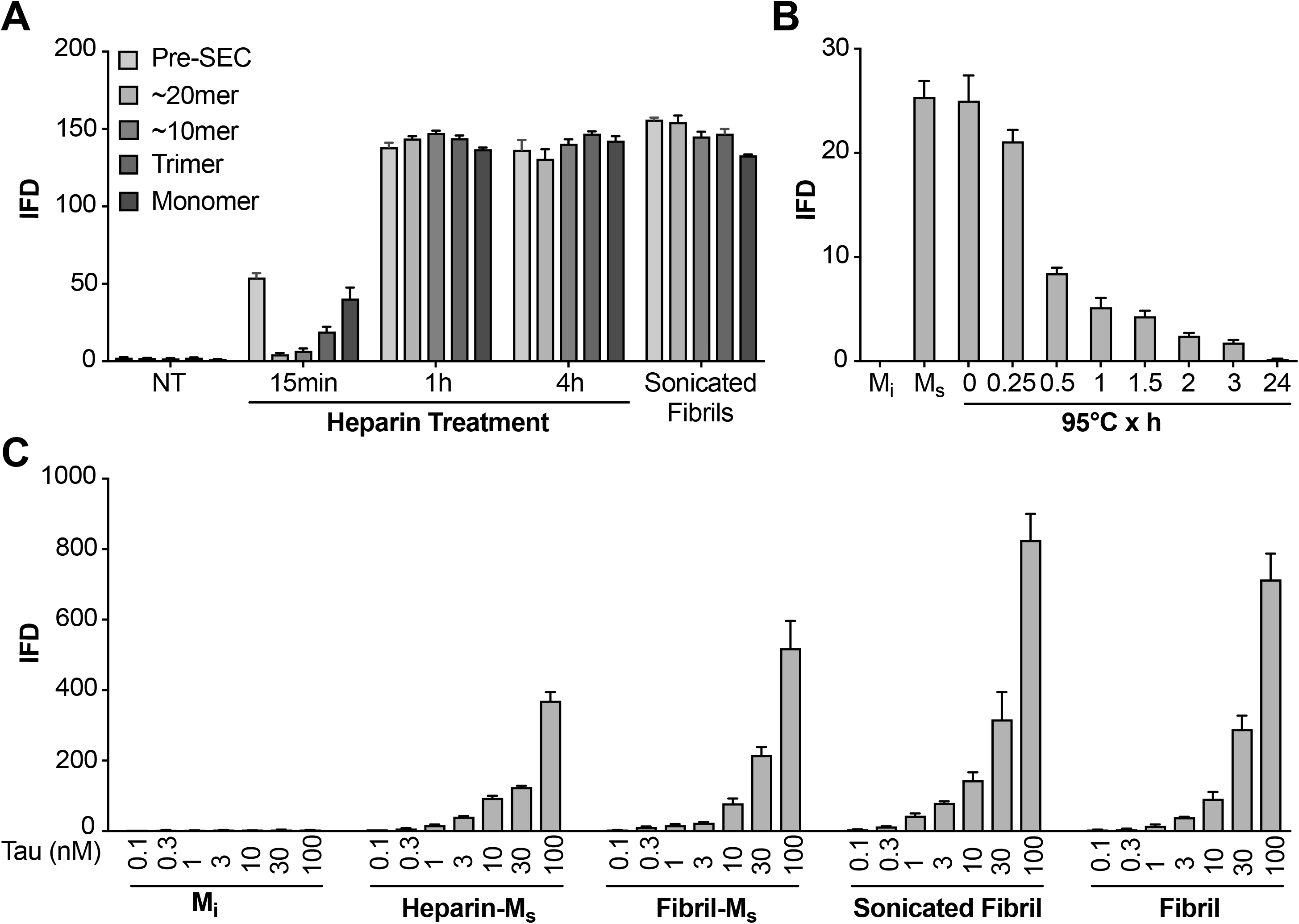
Heparin induces transition from M_i_ to M_s_. (**A**) Heparin treatment of FL WT tau was carried out for 15min, 1h, or 4h. Samples were resolved by SEC, and fractions of various sizes were compared using the biosensor seeding assay. “Pre-SEC” refers to the sample prior to fractionation. NT = monomer not treated with heparin. At 15min, a small, but significant seeding activity was observed primarily in the monomer fraction. By 1h this signal was very strong, and comparable to the signal of M_s_ derived from sonicated fibrils. (**B**) M_s_ derived from 4h heparin exposure was heated at 95°C for different times, followed by analysis of seeding activity in the FRET biosensor assay. Seeding activity decayed over 24h. (**C**) Seeding efficiencies per nM of tau (monomer equivalent) of the various forms of M_s_, sonicated, or unsonicated fibrils were relatively similar. M_i_ was sonicated identically to M_s_, followed by purification via SEC, but exhibited no seeding activity. Transfection of heparin failed to trigger intracellular aggregation (data not shown).

### XL-MS reveals unique contacts associated with M_s_

To probe the structures of M_i_ and M_s_, we employed cross-linking with mass spectrometry (XL-MS), which uses DSS-mediated crosslinking of proteins (monomer or larger assembly) followed by trypsin proteolysis, enrichment of resultant fragments by SEC, and identification by capillary liquid chromatography tandem mass spectrometry (MS). This method creates restraints for structural models of single proteins or protein complexes^23, 28, 29^. We assigned the complex fragment ion spectra to the corresponding peptide sequences using xQuest ^24^. Denaturation of recombinant tau with 8M urea prior to crosslinking produced no intramolecular cross-links (data not shown), indicating that crosslinks observed under native conditions represented local structure. We studied M_i_, fibril-derived M_s_ and heparin-derived M_s_ using XLMS. Short reaction times ensured the production of only intra-molecular crosslinks as monitored by SDS-PAGE (Fig. S6). XL-MS for each sample was carried out in triplicate (Suppl. Table S1), and only considering consensus crosslinks present in each replicate (Suppl. Table S2). M_i_ exhibited crosslink patterns which indicated local and distant intramolecular contacts (Fig. 7A). In M_s_, we observed a consistent crosslinking of K150 with K254, K267, K274 or K280 all located between RD 1 and 2. These crosslinks tracked exclusively with M_s_, both fibril- or heparin-derived (Fig. 7B,C). We never observed these crosslinks in M_i_. To test the relationship of this crosslink with seed function, we carried out heat denaturation at 95°C for 3 or 24h, followed by XL-MS. Heating samples results in a decrease in crosslink frequency (Fig. S7). Importantly, however, we observed a parallel persistence of this crosslink pattern with seeding activity (Fig. 7B,C). The XL-MS results indicate a distinct structure and seeding activity for M_s_ that is surprisingly resistant to denaturation at 95°C.

**Figure 7.**
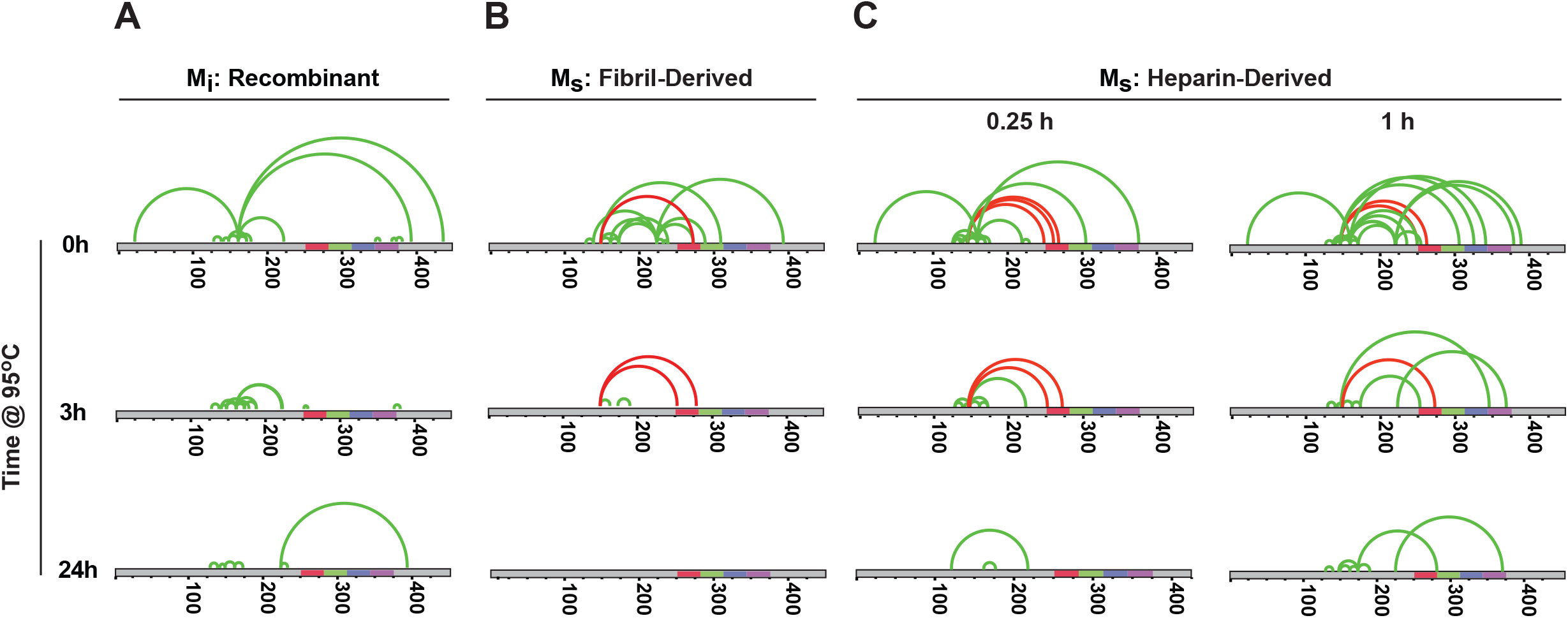
Unique XL-MS patterns for different forms of M_i_ and M_s_. Tau monomers were prepared as described, heated at 95°C for 0, 3 or 24h, reacted with DSS, proteolyzed and analyzed by mass spectrometry to define intramolecular crosslinks. Diagrams represent crosslinks within the tau protein. Tau is shown in grey; RD is colored in red (R1), green (R2), blue (R3) and indigo (R4). Each diagram indicates only crosslinks present in every triplicate (green or red). Crosslinks uniquely observed within M_s_ preparations are shown in red. Each sample was prepared, isolated by SEC, and then subjected XL-MS. (**A**) M_i_: tau monomer not previously fibrillized; (**B**) M_s_: fibril-derived tau monomer; (**C**) M_s_: heparin-exposed tau monomer (0.25h or 1h). Crosslinks from aa150 to aa254-290 mark all forms of M_s_ after exposure to 95°C for 0h, 0.25h and 3h, but not 24h.

### AD brain contains M_s_

Given our experiments with recombinant M_i_ and M_s_, we wished to test whether similar structures exist *in vivo*. We extracted AD and control brain samples using a dounce homogenizer to avoid liberating significant monomer from fibrils. We immunoprecipitated tau using an antibody that targets the amino-terminus (HJ8.5), and resolved the eluates by SEC, followed by ELISA to determine tau levels (Fig. 8A,B). Tau from control brain purified in the monomer fraction (Fig. 8A), while tau from AD brain distributed across multiple fractions, corresponding to monomer and larger assemblies (Fig. 8B). When we tested each fraction for seeding activity, we observed none in any control brain fraction (Fig. 8C). However, all AD fractions contained seeding activity, including monomer (Fig. 8C). To exclude the possibility that the brain homogenization protocol liberated M_s_ from neurofibrillary tangles, we spiked tau KO mouse brain samples with recombinant fibrils *in vitro*, or fibril-derived M_s_. We then used dounce homogenization and immuno-purification as for human brain. We evaluated the seeding activity in total lysate, supernatant following 10,000 x g centrifugation, and SEC fractions (Fig. 8D). We readily observed monomer seeding activity in tau KO brain spiked with M_s_, however we observed none in fractions that had been spiked with fibrils (Fig. 8D). The homogenization protocol for human brain was thus unlikely to have liberated M_s_ from pre-existing tau fibrils.

**Figure 8:**
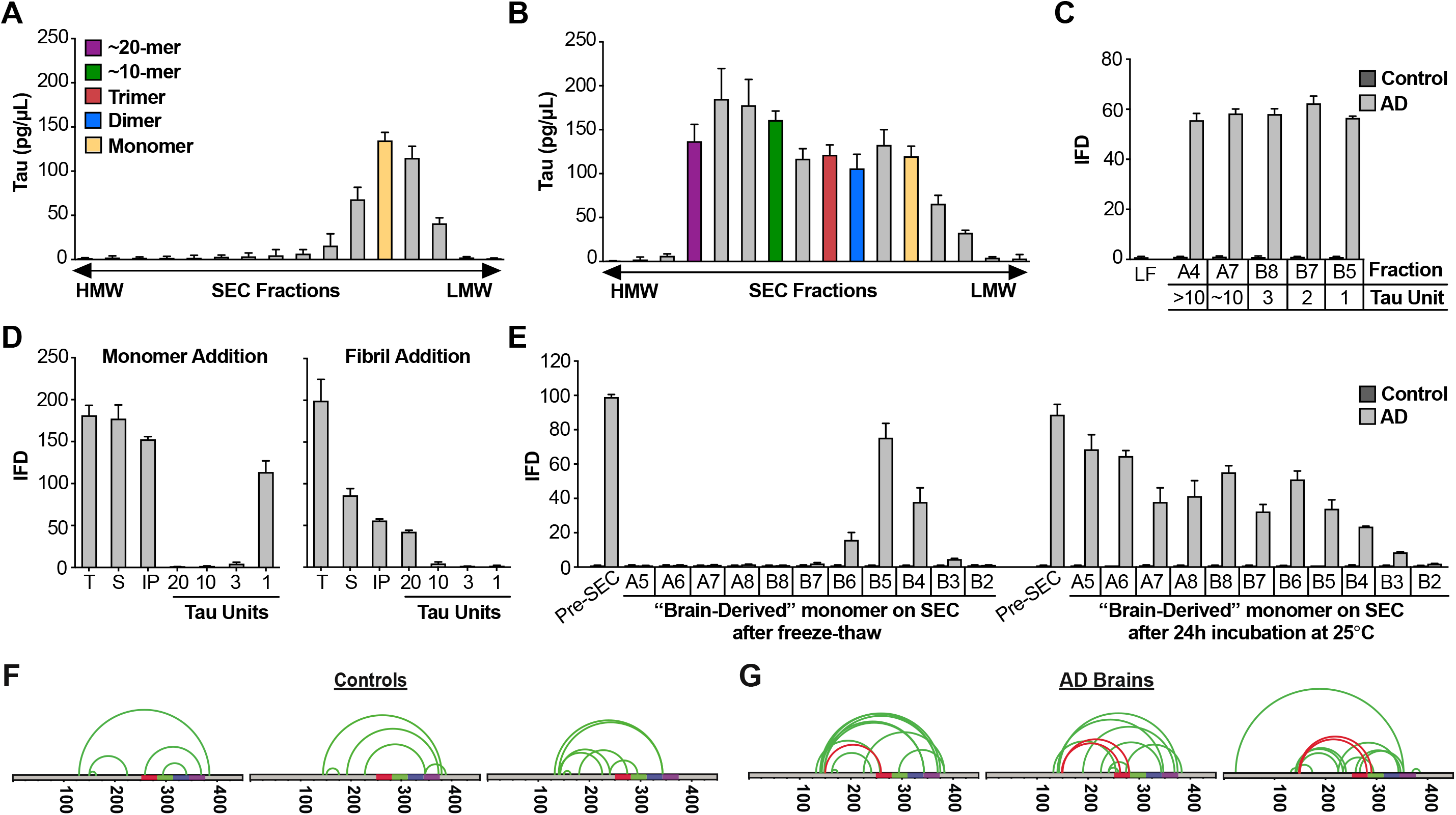
AD brain contains seed-competent monomer. Tau from control and AD brains was immunoprecipitated and subjected to SEC. (**A**) SEC from control brain contained predominantly tau monomer. (**B**) SEC from AD brain contained a range of tau assembly sizes. (**C**) Tau monomer from control brain exhibited no seeding activity, whereas monomer from AD brain did, along with larger assemblies. Tau Unit refers to the putative number of molecules per assembly. LF = Lipofectamine control. (**D**) Tau KO mouse brain was spiked either with human tau M_s_ or fibrils prior to dounce homogenization, immunopurification, and resolution by SEC. Samples spiked with M_s_ exhibited monomer seeding activity, but not samples that had been spiked with fibrils. (**E**) AD-derived tau monomer was incubated for the indicated times prior to SEC and determination of seeding activity in each fraction. Larger seed-competent assemblies formed after 24h incubation at RT. (**F, G**) Three control and AD brains were homogenized, monomer isolated, and evaluated by XL-MS. Tau monomer from controls lacked the long-range crosslinks observed in M_s_. AD-derived M_s_ contained long-range crosslinks (aa150 to aa254-290) also observed in recombinant forms of M_s_.

To test for self-association of control-derived M_i_ vs. AD-derived M_s_, we purified these species by SEC, and divided each monomer fraction in two. We snap-froze one fraction and incubated the other overnight at room temperature. Then we again resolved the assemblies via SEC and tested each fraction for seeding activity. Control monomer was inert, even after incubation at RT (Fig. 8E). AD-derived M_s_ that was purified, frozen, and re-purified by SEC exhibited seeding activity exclusively in the monomer fraction (Fig. 8E). By contrast, AD-derived M_s_ incubated at RT formed seed-competent assemblies of increasing size (Fig. 8E). We concluded that, as for other types of M_s_, AD-derived M_s_ exhibited an intrinsic capacity for self-association into seed-competent assemblies. To compare structures of control vs. AD-derived monomer via XL-MS, we isolated tau from brains of 3 AD patients and 3 age-matched controls. In control-derived monomer, we observed no evidence of the crosslink that marked M_s_ (Fig. 8G). However, in each AD-derived M_s_ sample we observed a discrete set of crosslinks between aa150 and aa259-290 (Fig. 8H). This essential finding did not change, no matter what method of homogenization we used (Supp. Fig. S8, Suppl. Table S3), and implied a common structure that unifies ensembles of seed-competent tau monomer, whether produced *in vitro* or *in vivo*.

### Models of seed-competent monomer suggest exposure of VQIINK and VQIVYK

Based on intramolecular FRET and electron paramagnetic resonance spin labeling Mandelkow et al. have previously proposed native tau structure to be in a “paperclip” configuration, with the C-terminus folded over the RD^30^. To understand how core elements of tau control its aggregation, we employed Rosetta to create models of tau structure for M_i_ and M_s_ using restraints from the crosslink patterns and length of the DSS crosslinker. The overall energetics and radii of gyration in the models were comparable for M_i_ and M_s_ (Fig. S9), indicating global structural similarity. We thus focused on the RD, given its high frequency of intramolecular crosslinks, and primary role in aggregation (Fig. 9A). We observed differences in the predicted interface structure between R1/R2 and R2/R3 which encode two core VQIINK and VQIVYK motifs critical for tau amyloid formation^31, 32^. The M_i_ structural model predicted masking of VQIINK and VQIVYK sequences in compact “hairpin” structures (Fig. 9B, Supp. Movie M_i_), similar to the structure of microtubule-bound tau previously determined by NMR ^33^. By contrast, within M_s_ the model predicted relative exposure of VQIINK and VQIVYK (Fig. 9C, Supp. Movie M_s_). We next evaluated XL-MS-guided predictions of patient-derived tau, although lower sample quality and fewer high confidence crosslinks (possibly due to protein heterogeneity) limited our accuracy. As for recombinant protein, M_i_ from control patients also featured VQIINK/VQIVYK sequences in a less accessible configuration (Fig. 9D, Supp. Table S1; Supp. Movie: Control1). In AD-derived M_s_, long-range contacts from aa150 to R2 influenced the model, and predicted an exposed configuration of VQIINK/VQIVYK (Fig. 9E, Table S1; Supp. Movie: AD1). With important caveats, the models guided by XL-MS imply that the general difference between M_i_ and M_s_ derives from relative shielding vs. exposure of VQIINK/VQIVYK sequences.

**Figure 9.**
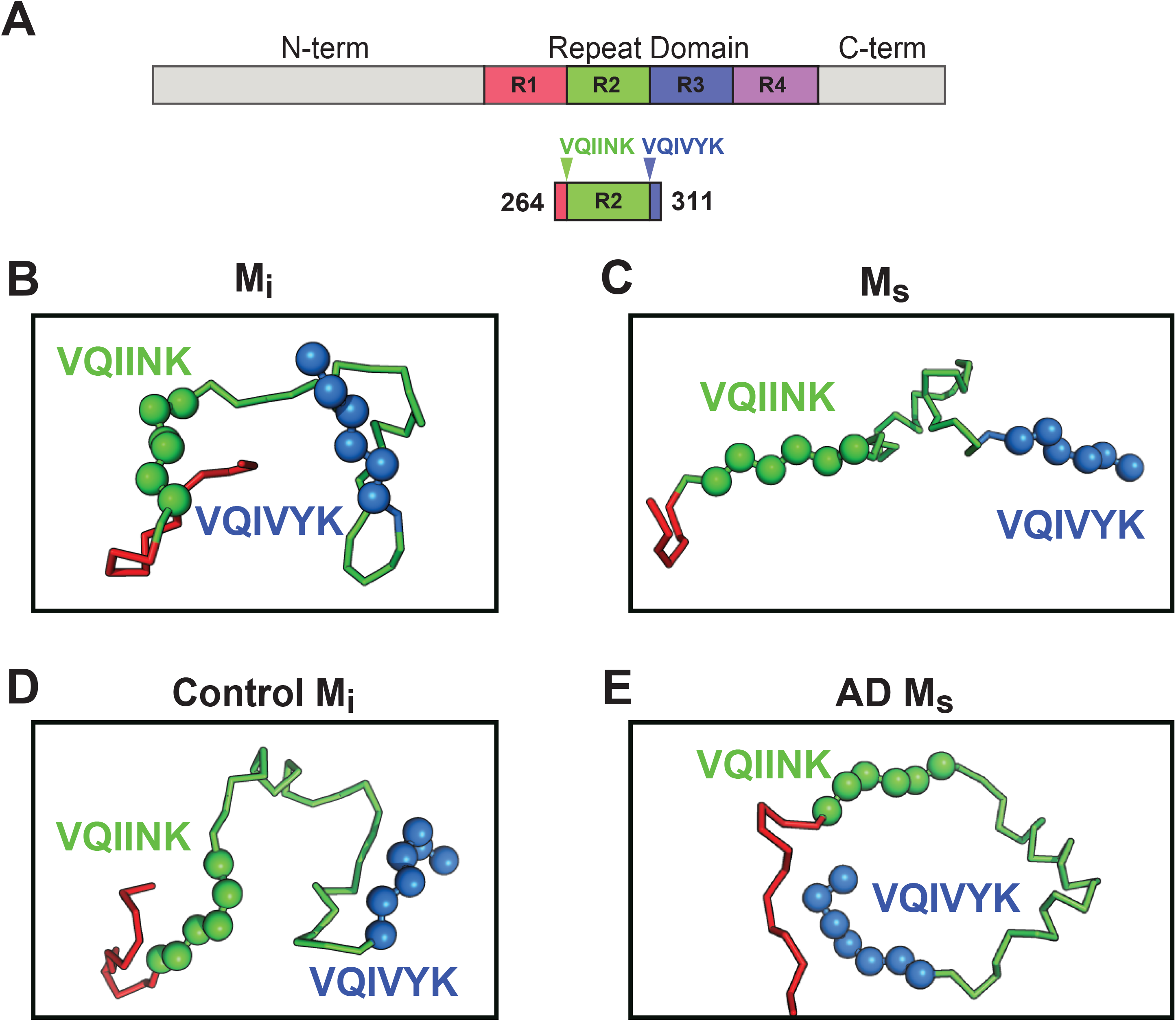
Models of M_i_ and M_s_ suggest differences in the R1R2 and R2R3 regions. XL-MS identified pairs were used as restraints in Rosetta to create structural models of discrete tau domains. (**A**) Schematic highlighting the region of the RD encoding structural differences between M_i_ and M_s_. Tau RD is colored in red (R1), green (R2), blue (R3) and indigo (R4); N-and C-terminal portions of tau are shown in grey. Fragments of interest are shown with their position in the RD. (**A**) recombinant M_i_; (**B**) fibril-derived M_s_, (**C**) Control M_i_ and (**D**) AD-derived M_s_. Regions surrounding the R1R2 and R2R3 are indicated, highlighting two amyloid-forming sequences, VQIINK (green spheres) and VQIVYK (blue spheres). In both forms of M_i_ VQIINK and VQIVYK are associated with flanking amino acids in hairpin structures. In both forms of M_s_ the VQIINK and VQIVYK sequences are presented at the protein surface. Please see Supplemental Movie files to better visualize the 3D orientation of specific regions.

### Limited proteolysis supports models of exposed VQIINK/VQIVYK sequences

As an orthogonal comparison of the structures of M_i_ and M_s_, we used limited proteolysis with trypsin. M_i_ or M_s_ (heparin-exposed) that had been passed through a 100kD filter immediately prior were subjected to a fine time course of limited proteolysis (Fig. 10A). Each sample was prepared in triplicate with matched protein quantities to facilitate label-free analysis. We then used mass spectrometry to evaluate the production of tau fragments and mapped these to specific cleavage sites. We identified 60 peptides common across the two conditions (Suppl. Table S4). To summarize enrichment of peptides across the two datasets we compared the ratio of averaged kinetic profiles (Fig. S10). Differences between the M_i_ and M_s_ primarily localized to the RD (Fig. S10). In M_i_, an R1R2 fragment was enriched (Fig. 10C) while only the R2 portion of that fragment was enriched in M_s_ (Fig. 10D). We observed similar patterns in R2R3 (Fig. 10F,G). By contrast, other domains outside of these regions had similar cleavage kinetics in M_i_ and M_s_ (Fig. 10E,H, Fig. S10). Mapping these cleavage sites onto our structural models revealed that proteolysis in M_i_ preferentially occurred outside the hairpin that includes VQIINK and VQIVYK amyloid sequences, while cleavage in M_s_ occurred adjacent to the amyloid sequences (Fig. 10I,J). The cleavage patterns were thus consistent with structural models of VQIINK and VQIVYK regions, which predicted relative inaccessibility of hairpin-associated sequences in M_i_, and accessibility in M_s_.

**Figure 10.**
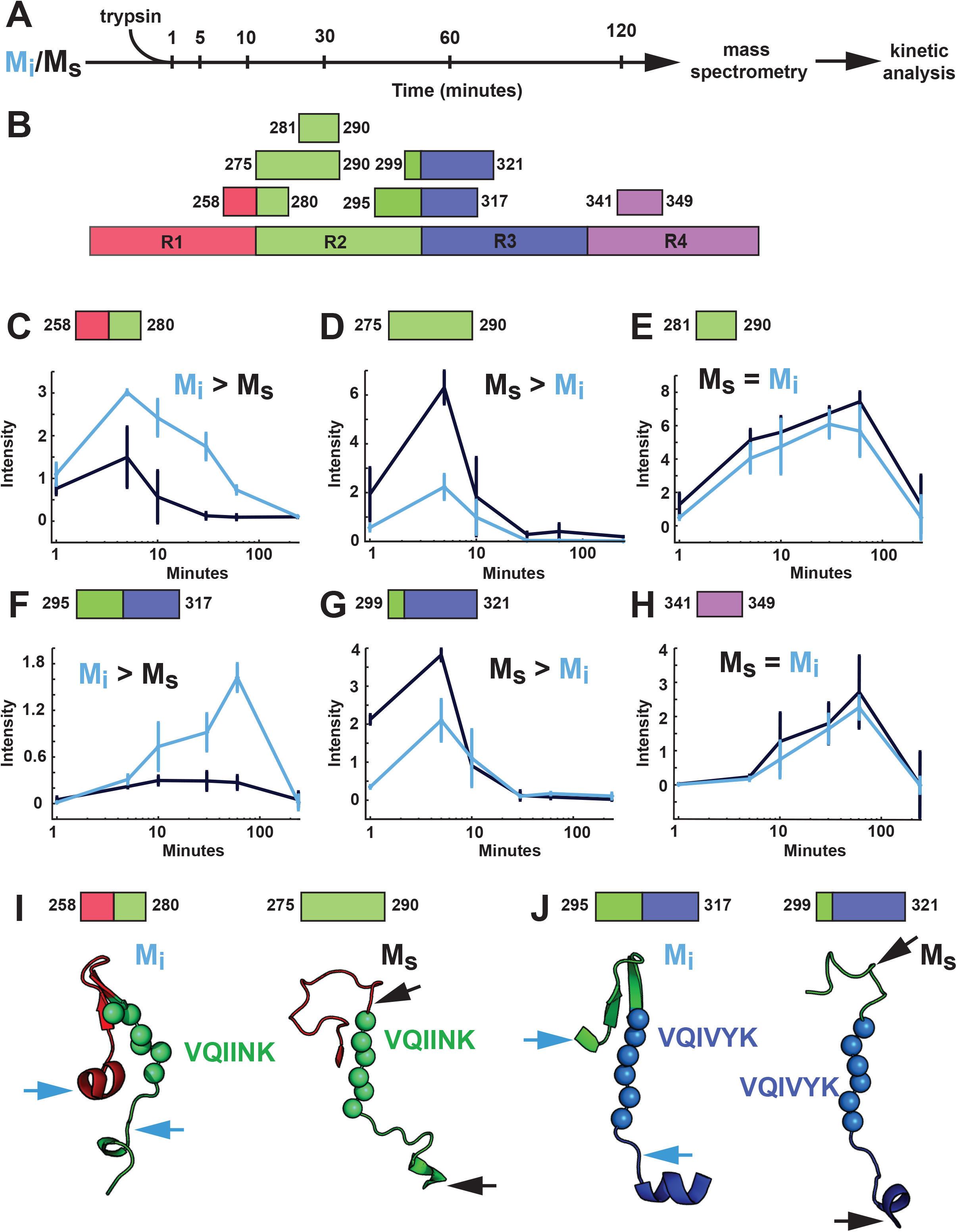
Proteolysis of M_i_ and M_s_ reveals distinct patterns. (**A**) M_i_ and M_s_ were prepared in triplicate, isolated by SEC, and passed through a 100kD filter immediately prior to exposure to trypsin for 1, 5, 10, 30, 60 and 120min. Samples were analyzed by mass spectrometry and kinetic profiles generated for peptides present at each time point. (**B**) Tau RD is colored in red (R1), green (R2), blue (R3) and indigo (R4). Identified peptides are shown with their position in the RD. (**C-H**) Kinetic profiles are indicated for peptides that were more abundant in M_i_ (**C, F**), M_s_ (**D, G**) or equal in M_i_ and M_s_ (**E, H**). M_i_ and M_s_ kinetic profiles are shown in blue and black, respectively. Fragments enriched in M_i_ or M_s_ were mapped onto corresponding regions in the structural models (**I, J**). The models are shown as cartoons colored in red (R1), green (R2) and blue (R3). Cleavage sites are indicated by arrows for M_i_ (blue) and M_s_ (black).

## Discussion

We propose that tau monomer occupies two distinct and stable conformational ensembles. One set of structures (collectively termed M_i_) is relatively inert, while another has intrinsic ability to self-assemble, and acts as a template, or seed, for fibril growth *in vitro* and in cells (collectively termed M_s_). Multiple controls indicated that our original preparation of fibril-derived M_s_ is in fact a monomer, uncontaminated by larger assemblies. Tau monomer purified from AD brain also had intrinsic seeding activity, and self-associated to produce larger seed-competent assemblies. A model restrained by the XL-MS data, and consistent with biochemical studies, predicts that VQIVYK and VQIINK sequences assume an open configuration in all types of M_s_ (fibril-derived, heparin-induced, and AD-derived). By contrast, the model predicts lack of VQIINK/VQIVYK exposure in M_i_. Limited proteolysis studies are consistent with this idea, although clearly more detailed biochemical, biophysical, and structural analyses will be needed to test its validity. Taken together, these data establish a new concept for tau: this intrinsically disordered protein has multiple, stable monomeric states, functionally distinguished by the presence or absence of seeding activity.

Amyloid proteins form progressively larger assemblies over time, and it has been difficult to define the composition of the minimal seed. Mandelkow and colleagues studied tau aggregation *in vitro* and concluded that a seed of 8-12 molecules existed in their experimental system^4^. By contrast, Kuret and colleagues posited an “intermediate” of tau that could subsequently initiate self-assembly, and their data, based on extrapolation of tau concentrations needed to enable development of thioflavin fluorescence *in vitro*, were consistent with a monomeric seed^1^. Wetzel and colleagues also proposed that a monomer is the basis of a “thermodynamic nucleus” that templates the aggregation of synthetic polyglutamine peptides^34^. However, no prior study has previously identified stable forms of tau monomer that seed amyloid formation.

The actual cause of tau aggregation in tauopathies is unknown. It has been proposed that dissociation of tau monomer from microtubules, possibly due to phosphorylation, allows high concentration and self-association to form pathogenic assemblies^35^. In this study, using a single source of recombinant protein, we define distinctly structured seed-competent and inert forms of tau. We have similarly identified seed-competent species in human brain. In reality “seed-competent” and “inert” forms of tau almost certainly represent multiple structural ensembles separated by defined energy and/or kinetic barriers. The barrier for conversion of an inert to a seed-competent form of tau can apparently be overcome by incubation with heparin and/or incorporation into a fibril. In neurons, other factors such as post-translational modifications and heterologous binding events likely play a role. Identification of the factors that trigger conversion from inert to seed-competent forms will thus have obvious implications for understanding disease mechanisms.

Isolation of seed-competent monomer from AD brain, with a very mild purification that explicitly excludes sonication or vigorous tissue homogenization, strongly suggests that this form of tau exists *in vivo*. Furthermore, we observed that both recombinant M_s_ and AD-derived M_s_ build multimeric assemblies *in vitro* far more efficiently than M_i_ or control-derived monomer. Thus, we hypothesize that a uniquely structured form of tau may be required for efficient assembly growth in cells. This contrasts with the idea that multimeric assemblies uniquely stabilize the conformation of otherwise unstructured proteins as they incorporate into the growing fibril, or that liquid-liquid phase separation with extremely high local concentration underlies tau aggregation^36^. Instead, we imagine that the initiation of aggregation in human brain might begin with a stable transition of tau monomer from an inert to a seed-competent form. To fully study this process will require more extensive biochemical purification of tau M_s_ from the earliest stages of disease.

M_s_ has a remarkably stable structure, as it resists heat denaturation at 95°C for up to 3h. This suggests a heretofore unrecognized conformation of tau that, to account for its slow denaturation, likely involves multiple intra-molecular interactions involving short and long range amino acid contacts. XL-MS provides some indication of what these might be, and crosslinks between aa150 and R2 appear to mark a seed-competent conformation. In agreement with the XL-MS results, we observed that heat inactivation of M_s_ seeding activity occurs with a lag phase, rather than first order time-dependent decay. This implies a complex tertiary structure in which M_s_ has multiple seed-competent intermediates. Future XL-MS studies performed at different temperatures could reveal these structures. With more advanced methods to interrogate the structure of monomeric tau in patient material, we imagine that “seed-competent monomer” will in fact represent myriad structures, depending on the underlying disease. This could provide an explanation for how a single tau protein might self-assemble into diverse amyloid strains. We note with excitement a recent study of the yeast prion Sup35 from the Tanaka laboratory. Like tau, Sup35 is intrinsically disordered, yet they have observed local structure that influences the conformations of fibrils it can form^37^.

Without further studies to identify structures of tau at higher resolution, we cannot know for certain why one form acts as a seed and another does not. However, we gained important insights when we modeled the configurations of R1R2 and R2R3 using Rosetta, with crosslinks as restraints. With obvious caveats, our models predicted that the local environment surrounding two hexapeptide motifs, VQIINK and VQIVYK, which are required for tau to form amyloid structures, may explain the differences between seed-competent and inert forms. In the models of M_i_, and control brain-derived tau, these motifs lie buried in hairpin structures. By contrast, in M_s_ and AD-derived tau, both are exposed. VQIINK and VQIVYK thus might mediate intermolecular interaction in a growing assembly. In support of our structural model, the proteolysis experiments corroborate differences in exposure of the VQIINK and VQIVYK sequences in the R1R2 and R2R3 regions between M_i_ and M_s_. We note with great enthusiasm the recent study of Fitzpatrick et al. ^38^, which defined critical sequences of tau within the amyloid core that are based on VQIVYK and adjacent amino acids. Indeed, it has been recently observed that heparin binding involves residues spanning 270-290, and promotes expansion of the remainder of the molecule ^39^. This is consistent with our predictions of relative exposure of VQIINK/VQIVYK. The diversity of exposed core elements (almost certainly beyond VQIINK/VQIVYK) could specify the formation of assemblies that give rise to distinct strains, as suggested by work from the Tanaka laboratory^37^. Consistent with this idea, the Fitzpatrick et al. study indicates that in AD-derived tau fibrils the VQIVYK sequence plays a key role in the core amyloid structure (along with adjacent amino acids), but the VQIINK sequence does not ^38^. We also note that multiple disease-associated mutations in tau affect residues in close proximity to VQIINK/VQIVYK. For example, our models predict that serine or leucine substitutions at P301 (which cause dominantly inherited tauopathy) would uniquely destabilize the local structure and promote exposure of the VQIINK/VQIVYK sequences. Future experiments will test these ideas more definitively.

## Acknowledgements

We thank Peter Davies for generously providing antibody reagents and ELISA protocol guidance. This work was supported by grants from the Tau Consortium and NIH grants awarded to 1R01NS071835 (M.I.D.), R01NS089932 (R.V.P. and M.I.D.), and the Effie Marie Cain Endowed Scholarship (L.A.J.). We appreciate the help of the Live Cell Imaging Core Facility administered by Katherine Luby-Phelps, Ph.D., and the Proteomics Core Facility at the University of Texas Southwestern Medical Center.

## Competing Interests

A patent disclosure has been filed by H.M., L.A.J. and M.I.D. related to the use of unique crosslinks to create biomarkers for neurodegenerative diseases.

---

**Supplemental Table S1. Summary of triplicate XLMS datasets**

**Supplemental Table S2. Summary of consensus XLMS datasets**

**Supplemental Table S3. Summary of patient-derived XLMS datasets**

**Supplemental Table S4. Summary of peptides identified in the M_i_ and M_s_ proteolysis**

## Supplemental Movie Files

PyMol was used to create rotating movies of all structural models for M_i_ and M_s_ derived from recombinant or human sources. Each model of M_s_ features one or both VQIINK/VQIVYK sequences exposed. Forms of M_i_ feature these sequences relatively buried in hairpin structures.

---

**Supplemental Figure S6.**
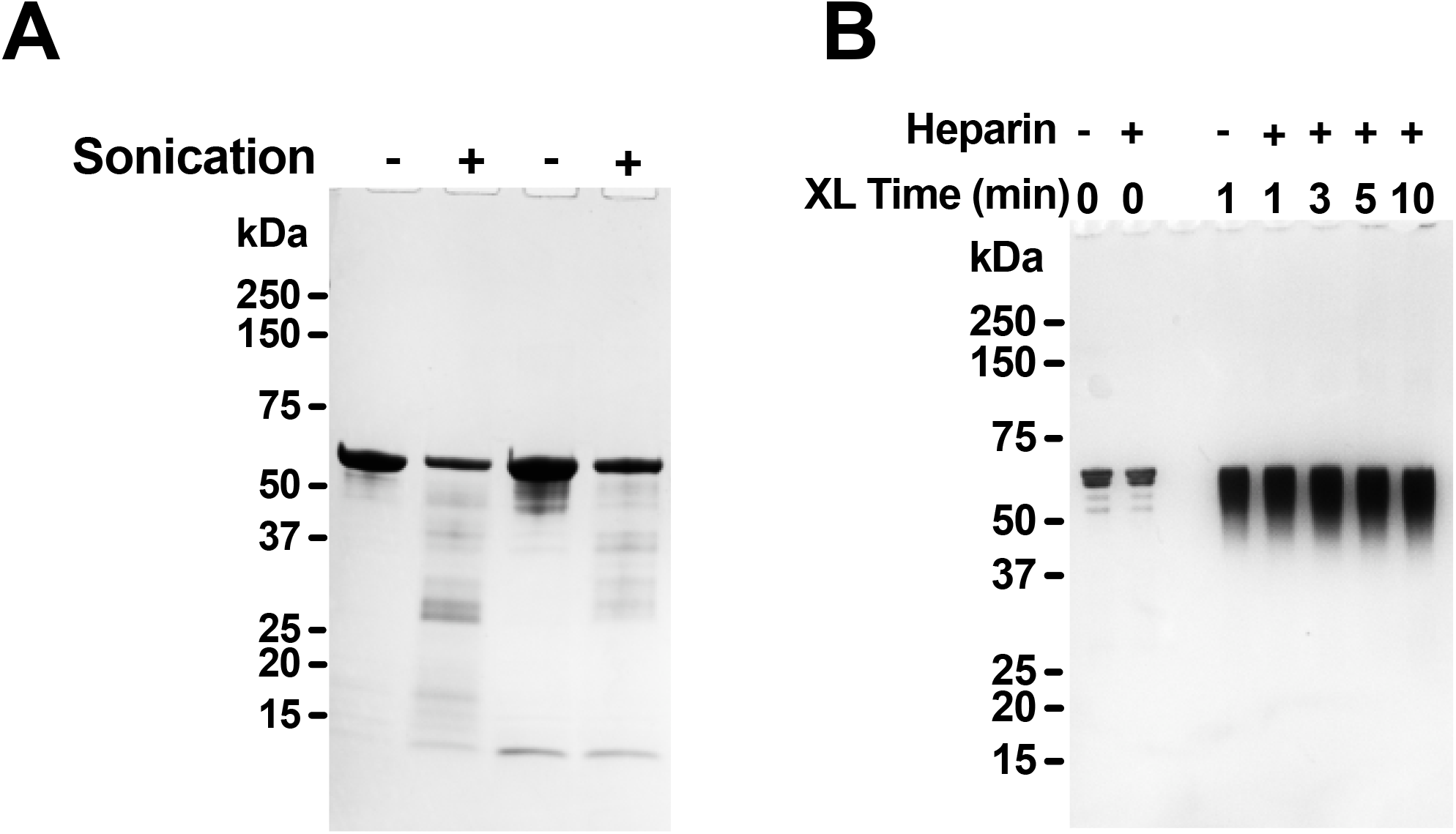
SDS-PAGE of tau after sonication or heparin treatment. (**A**) Two different FL WT tau preparations were sonicated or not, and 1.5µg protein was then resolved by SDS-PAGE and coomassie stain. Sonication induced a small degree of protein fragmentation. (**B**) FL WT tau was exposed to heparin for 15min, sufficient to induce conversion from M_i_ to M_s_, followed by DSS crosslinking for the indicated time periods. 100ng Protein was then resolved by SDS-PAGE and silver stain. No small fragments or higher-order crosslinked species were visible.

**Supplemental Figure S7.**
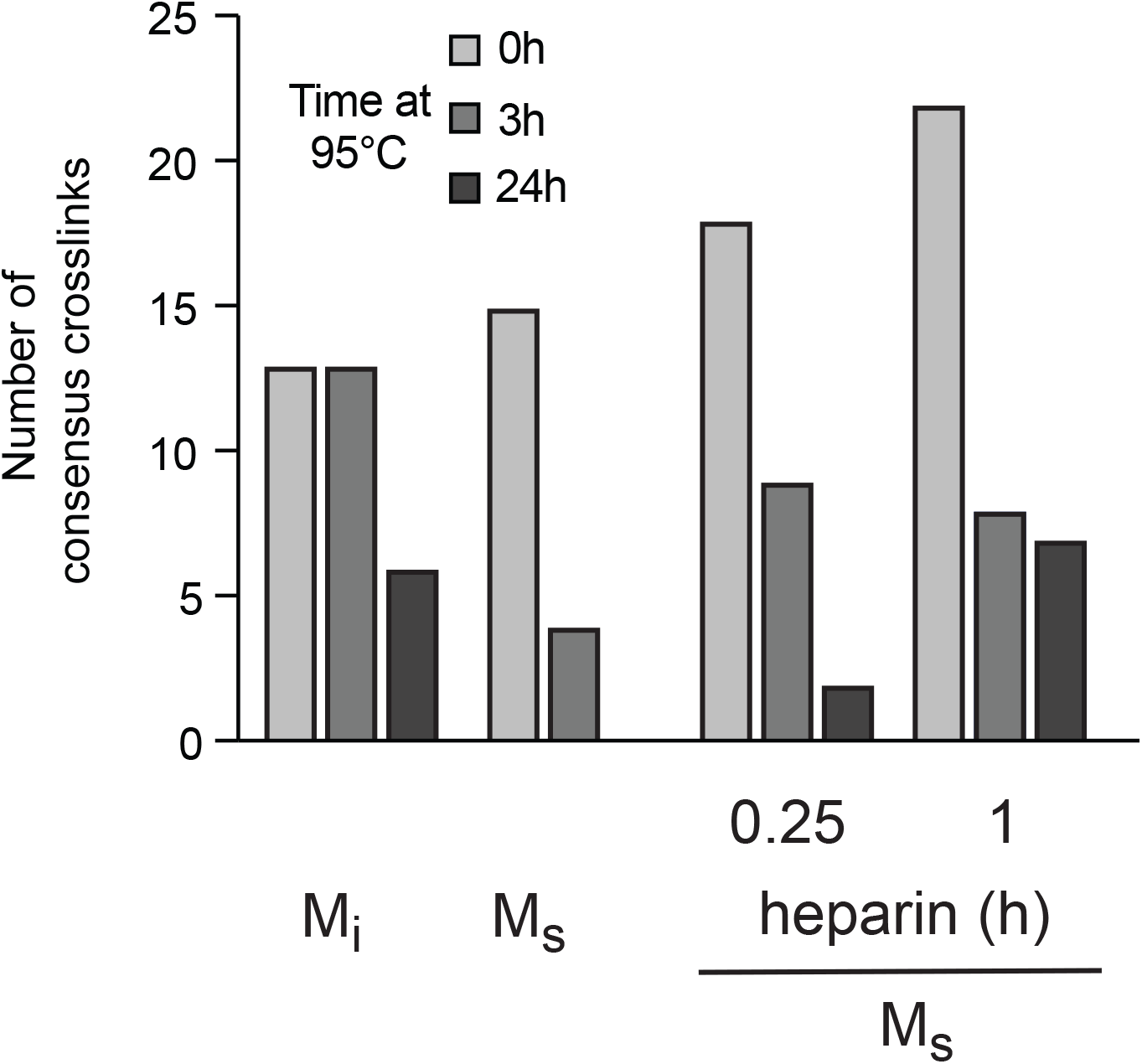
Frequency of crosslinks decrease with heat incubation. Heat denaturation of M_i_ and M_s_ (fibril-derived and heparin treated for 0.25h, 1h) decreases the abundance of consensus crosslink pairs (**A**). Columns represent data after exposure to 95°C for 0h, 3h and 24h.

**Supplemental Figure S8.**
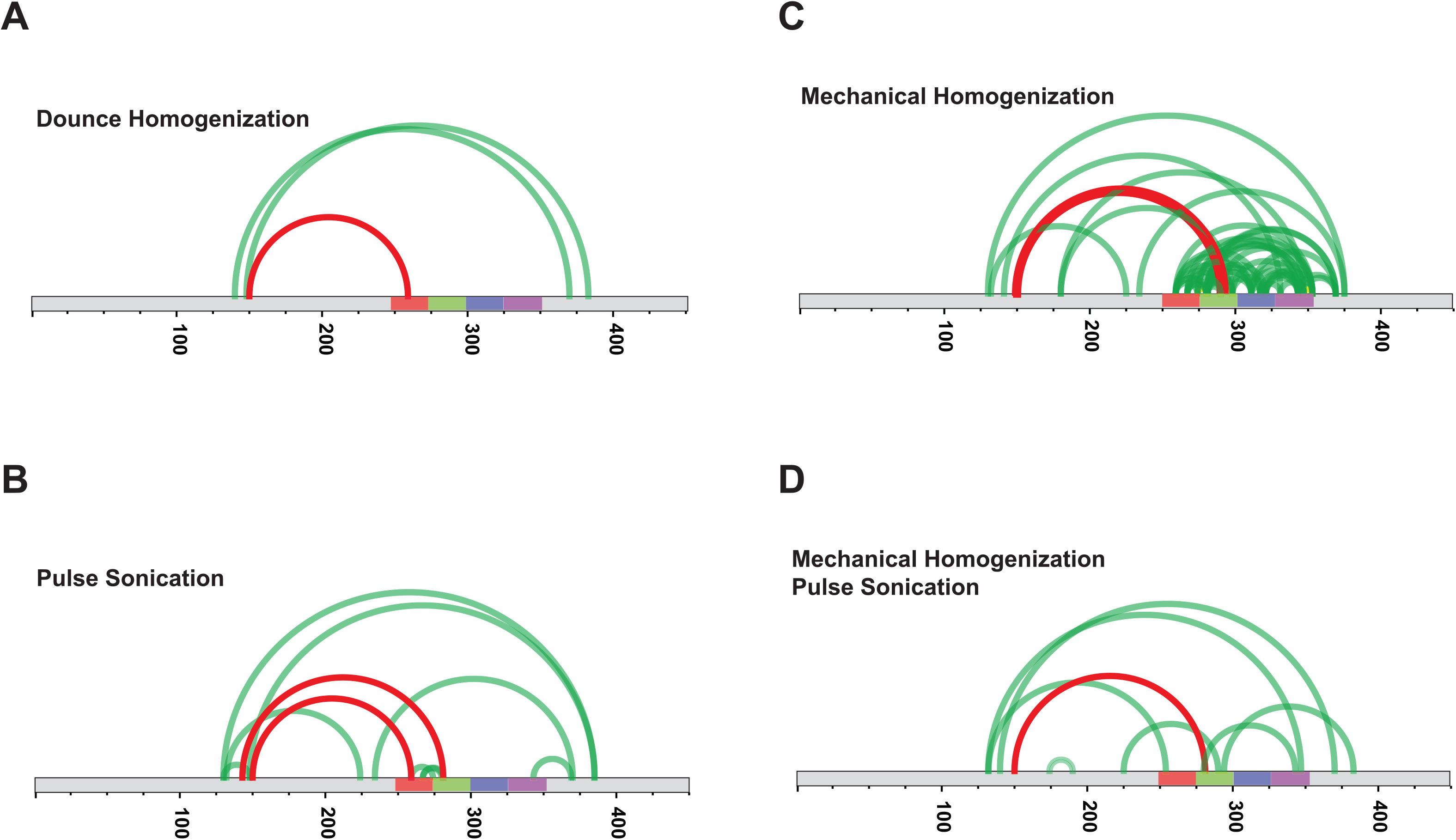
Different brain homogenization methods yield similar crosslink patterns. A single AD brain sample was homogenized using four different methods: (**A**) Dounce homogenization; (**B**) Pulse sonication; (**C**) Mechanical homogenization; (**D**) Mechanical homogenization followed by pulse sonication. Diagrams represent crosslinks within the FL tau protein. RD is colored in red (R1), green (R2), blue (R3) and indigo (R4). High confidence XL-MS crosslinks are shown as light green lines; crosslinks found in M_s_ are shown in red.

**Supplemental Figure S9.**
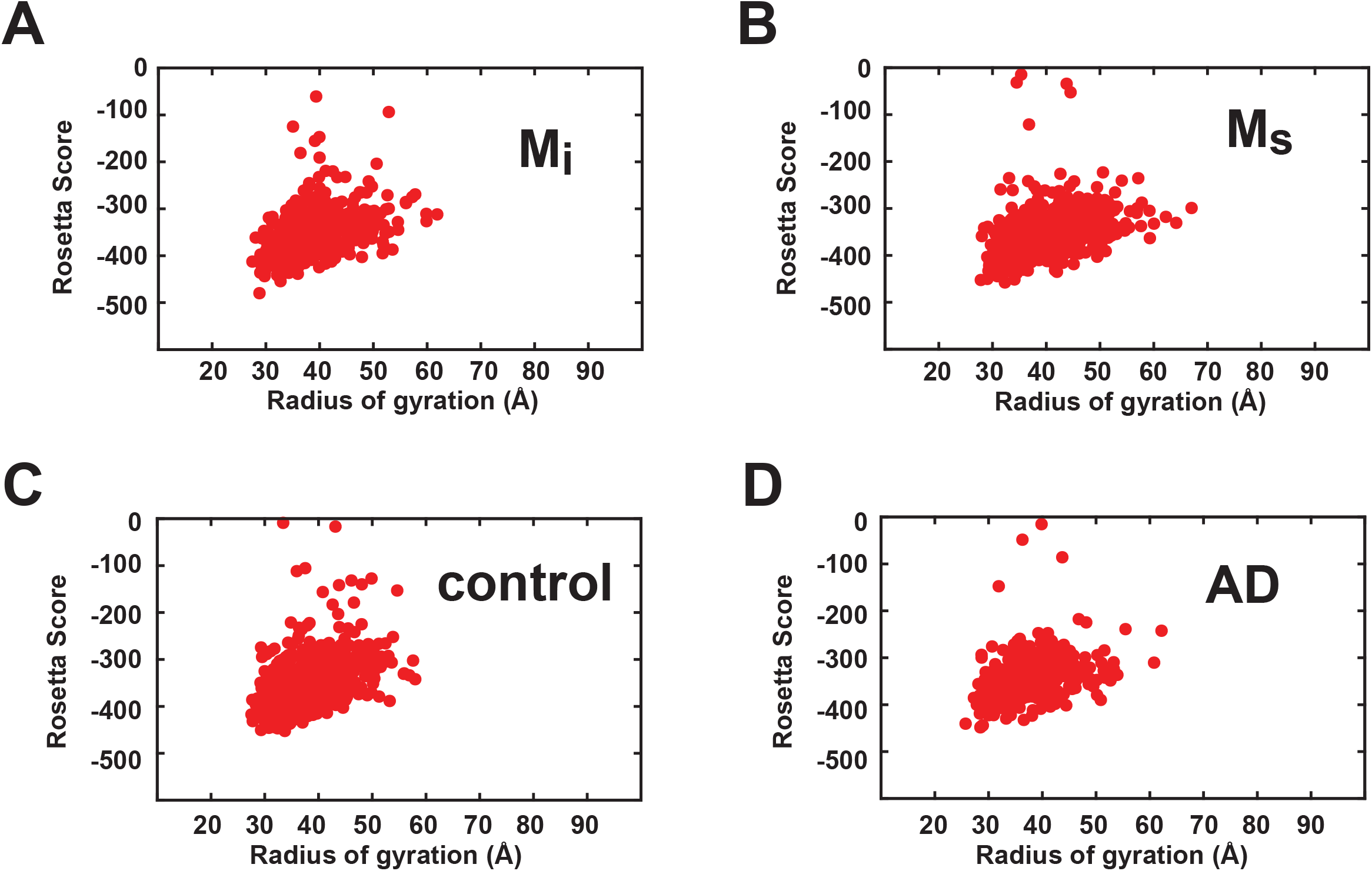
Energetics of Rosetta structural ensembles. The ensembles are shown as a distribution of total energy of each model and radius of gyration for recombinant M_i_ (**A**), recombinant M_s_ (**B**), control brain-derived M_i_ (**C**) and AD-derived M_s_ (**D**).

**Supplemental Figure S10.**
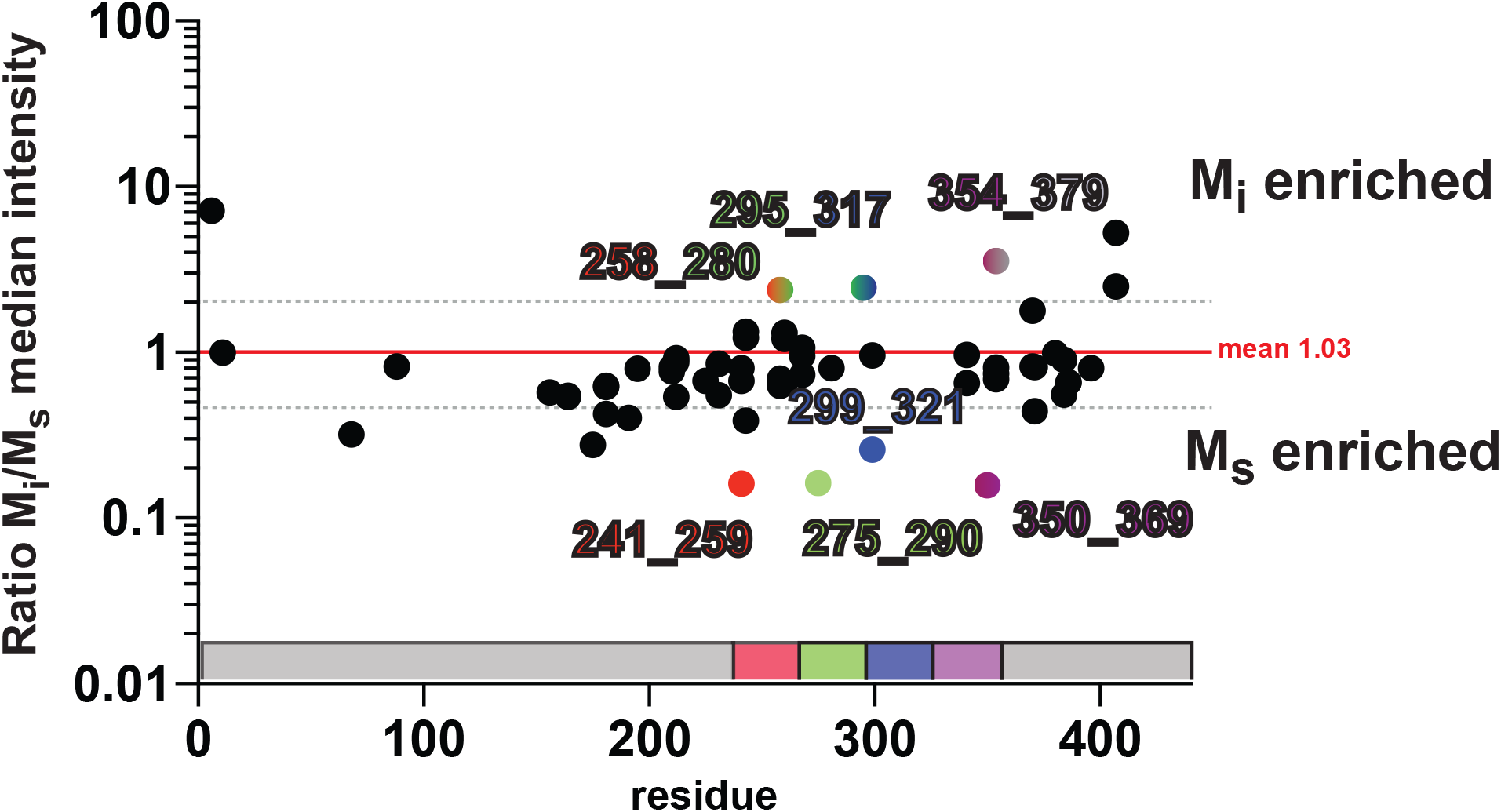
Proteolysis reveals localized differences between M_i_ and M_s_. The medians of the averaged kinetic profiles were compared as ratios for M_i_ and M_s_. The data were compared to the mean (red line) and standard deviation (dotted grey line). Peptides within the RD that are enriched in M_i_ or M_s_ are shown as colored dots according to location in the RD and labeled with N-term and C-term peptide positions. As a reference the tau RD is colored in red (R1), green (R2), blue (R3) and indigo (R4). Identified peptides are shown with their position in the RD.

## References

1. Chirita CN, Congdon EE, Yin H, Kuret J. Triggers of full-length tau aggregation: a role for partially folded intermediates. Biochemistry. 2005 Apr 19;44(15):5862–72.

2. Kar K, Jayaraman M, Sahoo B, Kodali R, Wetzel R. Critical nucleus size for disease-related polyglutamine aggregation is repeat-length dependent. Nat. Struct. Mol. Biol. 2011 Mar;18(3):328–36. PMCID: PMC3075957

3. Ramachandran G, Udgaonkar JB. Mechanistic studies unravel the complexity inherent in tau aggregation leading to Alzheimer’s disease and the tauopathies. Biochemistry. 2013 Jun 18;52(24):4107–26.

4. Friedhoff P, Bergen von M, Mandelkow EM, Davies P, Mandelkow E. A nucleated assembly mechanism of Alzheimer paired helical filaments. Proc. Natl. Acad. Sci. U.S.A. National Academy of Sciences; 1998 Dec 22;95(26):15712–7. PMCID: PMC28109

5. Neurodegenerative tauopathies. 2001;24(1):1121–59. Retrieved from: http://www.annualreviews.org/doi/abs/10.1146/annurev.neuro.24.1.1121

6. Sanders DW, Kaufman SK, Holmes BB, Diamond MI. Prions and Protein Assemblies that Convey Biological Information in Health and Disease. Neuron. Elsevier; 2016 Feb 3;89(3):433–48. PMCID: PMC4748384

7. Tau Trimers Are the Minimal Propagation Unit Spontaneously Internalized to Seed Intracellular Aggregation. 2015 Jun 12;290(24):14893–903. PMCID: PMC4463437

8. Frost B, Jacks RL, Diamond MI. Propagation of tau misfolding from the outside to the inside of a cell. J. Biol. Chem. American Society for Biochemistry and Molecular Biology; 2009 May 8;284(19):12845–52. PMCID: PMC2676015

9. Holmes BB, Furman JL, Mahan TE, Yamasaki TR, Mirbaha H, Eades WC, et al. Proteopathic tau seeding predicts tauopathy in vivo. Proc. Natl. Acad. Sci. U.S.A. 2014 Oct 14;111(41):E4376–85. PMCID: PMC4205609

10. Holmes BB, Devos SL, Kfoury N, Li M, Jacks R, Yanamandra K, et al. Heparan sulfate proteoglycans mediate internalization and propagation of specific proteopathic seeds. Proc. Natl. Acad. Sci. U.S.A. National Acad Sciences; 2013 Aug 13;110(33):E3138–47. PMCID: PMC3746848

11. Sanders DW, Kaufman SK, Devos SL, Sharma AM, Mirbaha H, Li A, et al. Distinct Tau Prion Strains Propagate in Cells and Mice and Define Different Tauopathies. Neuron. 2014 May 21. PMCID: PMC4171396

12. Furman JL, Holmes BB, Diamond MI. Sensitive Detection of Proteopathic Seeding Activity with FRET Flow Cytometry. J Vis Exp. 2015;(106):e53205–5. PMCID: PMC4692784

13. Goedert M, Jakes R, Spillantini MG, Hasegawa M, Smith MJ, Crowther RA. Assembly of microtubule-associated protein tau into Alzheimer-like filaments induced by sulphated glycosaminoglycans. Nature. Nature Publishing Group; 1996 Oct 10;383(6600):550–3.

14. Pérez M, Valpuesta JM, Medina M, Montejo de Garcini E, Avila J. Polymerization of tau into filaments in the presence of heparin: the minimal sequence required for tau-tau interaction. J. Neurochem. 1996 Sep;67(3):1183–90.

15. Frost B, Ollesch J, Wille H, Diamond MI. Conformational diversity of wild-type Tau fibrils specified by templated conformation change. J. Biol. Chem. American Society for Biochemistry and Molecular Biology; 2009 Feb 6;284(6):3546–51. PMCID: PMC2635036

16. Acker CM, Forest SK, Zinkowski R, Davies P, d’Abramo C. Sensitive quantitative assays for tau and phospho-tau in transgenic mouse models. Neurobiol. Aging. 2013 Jan;34(1):338–50. PMCID: PMC3474864

17. Measurement of microsecond dynamic motion in the intestinal fatty acid binding protein by using fluorescence correlation spectroscopy. 2002 Oct 29;99(22):14171–6. Retrieved from: http://eutils.ncbi.nlm.nih.gov/entrez/eutils/elink.fcgi?dbfrom=pubmed&id=12381795&retmode=ref&cmd=prlinks

18. Spectroscopic Study and Evaluation of Red-Absorbing Fluorescent Dyes. 2003 Jan;14(1):195–204. Retrieved from: http://pubs.acs.org/doi/abs/10.1021/bc025600x

19. Morozova OA, March ZM, Robinson AS, Colby DW. Conformational features of tau fibrils from Alzheimer’s disease brain are faithfully propagated by unmodified recombinant protein. Biochemistry. American Chemical Society; 2013 Oct 8;52(40):6960–7. PMCID: PMC4142060

20. Yanamandra K, Kfoury N, Jiang H, Mahan TE, Ma S, Maloney SE, et al. Anti-Tau Antibodies that Block Tau Aggregate Seeding In Vitro Markedly Decrease Pathology and Improve Cognition In Vivo. Neuron. 2013 Oct 16;80(2):402–14. PMCID: PMC3924573

21. Laidler KJ. The development of the Arrhenius equation. Journal of Chemical Education. 1984.

22. Burnham KP, Anderson DR. Model selection and multimodal inference: a practical information-theoretic approach. p. 61–3.

23. Leitner A, Joachimiak LA, Bracher A, Mönkemeyer L, Walzthoeni T, Chen B, et al. The Molecular Architecture of the Eukaryotic Chaperonin TRiC/CCT. Structure. 2012 May;20(5):814–25.

24. Rinner O, Seebacher J, Walzthoeni T, Mueller LN, Beck M, Schmidt A, et al. Identification of cross-linked peptides from large sequence databases. Nat. Methods. Nature Publishing Group; 2008 Apr;5(4):315–8. PMCID: PMC2719781

25. Grimm M, Zimniak T, Kahraman A. xVis: a web server for the schematic visualization and interpretation of crosslink-derived spatial restraints. Nucleic acids …. 2015.

26. Kahraman A, Herzog F, Leitner A, Rosenberger G, Aebersold R, Malmström L. Cross-link guided molecular modeling with ROSETTA. Fernandez-Fuentes N, editor. PLoS ONE. Public Library of Science; 2013;8(9):e73411. PMCID: PMC3775805

27. Lange OF, Rossi P, Sgourakis NG. Determination of solution structures of proteins up to 40 kDa using CS-Rosetta with sparse NMR data from deuterated samples. 2012.

28. Lasker K, Förster F, Bohn S, Walzthoeni T, Villa E, Unverdorben P, et al. Molecular architecture of the 26S proteasome holocomplex determined by an integrative approach. Proc. Natl. Acad. Sci. U.S.A. National Acad Sciences; 2012 Jan 31;109(5):1380–7. PMCID: PMC3277140

29. Joachimiak LA, Walzthoeni T, Liu CW, Aebersold R, Frydman J. The structural basis of substrate recognition by the eukaryotic chaperonin TRiC/CCT. Cell. 2014 Nov 20;159(5):1042–55. PMCID: PMC4298165

30. Jeganathan S, Bergen von M, Brutlach H, Steinhoff H-J, Mandelkow E. Global hairpin folding of tau in solution. Biochemistry. 2006 Feb 21;45(7):2283–93.

31. Bergen von M, Friedhoff P, Biernat J, Heberle J, Mandelkow EM, Mandelkow E. Assembly of tau protein into Alzheimer paired helical filaments depends on a local sequence motif ((306)VQIVYK(311)) forming beta structure. Proc. Natl. Acad. Sci. U.S.A. National Academy of Sciences; 2000 May 9;97(10):5129–34. PMCID: PMC25793

32. Bergen von M, Barghorn S, Li L, Marx A, Biernat J, Mandelkow EM, et al. Mutations of tau protein in frontotemporal dementia promote aggregation of paired helical filaments by enhancing local beta-structure. J. Biol. Chem. 2001 Dec 21;276(51):48165–74.

33. Kadavath H, Jaremko M, Jaremko Ł, Biernat J, Mandelkow E, Zweckstetter M. Folding of the Tau Protein on Microtubules. Angew. Chem. Int. Ed. Engl. WILEY-VCH Verlag; 2015 Aug 24;54(35):10347–51.

34. Bhattacharyya AM, Thakur AK, Wetzel R. polyglutamine aggregation nucleation: thermodynamics of a highly unfavorable protein folding reaction. Proc. Natl. Acad. Sci. U.S.A. 2005 Oct 25;102(43):15400–5. PMCID: PMC1266079

35. Mandelkow E-M, Mandelkow E. Biochemistry and cell biology of tau protein in neurofibrillary degeneration. Cold Spring Harb Perspect Med. 2012 Jul;2(7):a006247–7. PMCID: PMC3385935

36. Wegmann S, Eftekharzadeh B, Tepper K, Zoltowska KM, Bennett RE, Dujardin S, et al. Tau protein liquid-liquid phase separation can initiate tau aggregation. EMBO J. 2018 Feb 22;:e98049.

37. Ohhashi Y, Yamaguchi Y, Kurahashi H, Kamatari YO, Sugiyama S, Uluca B, et al. Molecular basis for diversification of yeast prion strain conformation. Proc. Natl. Acad. Sci. U.S.A. National Academy of Sciences; 2018 Mar 6;115(10):2389–94.

38. Fitzpatrick AWP, Falcon B, He S, Murzin AG, Murshudov G, Garringer HJ, et al. Cryo-EM structures of tau filaments from Alzheimer’s disease. Nature. 2017 Jul 5;56:343. PMCID: PMC5552202

39. Zhao J, Huvent I, Lippens G, Eliezer D, Zhang A, Li Q, et al. Glycan Determinants of Heparin-Tau Interaction. Biophys. J. 2017 Mar 14;112(5):921–32. PMCID: PMC5355497

